# Rapid profiling of protein complex re-organization in perturbed systems

**DOI:** 10.1101/2021.12.17.473177

**Authors:** Isabell Bludau, Charlotte Nicod, Claudia Martelli, Peng Xue, Moritz Heusel, Andrea Fossati, Federico Uliana, Fabian Frommelt, Ruedi Aebersold, Ben C. Collins

**Author notes:** equal contribution (Isabell Bludau, Charlotte Nicod, Claudia Martelli).

## Abstract

Protein complexes constitute the primary functional modules of cellular activity. To respond to perturbations, complexes undergo changes in their abundance, subunit composition or state of modification. Understanding the function of biological systems requires global strategies to capture this contextual state information on protein complexes and interaction networks. Methods based on co-fractionation paired with mass spectrometry have demonstrated the capability for deep biological insight but the scope of studies using this approach has been limited by the large measurement time per biological sample and challenges with data analysis. As such, there has been little uptake of this strategy beyond a few expert labs into the broader life science community despite rich biological information content. We present a rapid integrated experimental and computational workflow to assess the re-organization of protein complexes across multiple cellular states. It enables complex experimental designs requiring increased sample/condition numbers. The workflow combines short gradient chromatography and DIA/SWATH mass spectrometry with a data analysis toolset to quantify changes in complex organization. We applied the workflow to study the global protein complex rearrangements of THP-1 cells undergoing monocyte to macrophage differentiation and a subsequent stimulation of macrophage cells with lipopolysaccharide. We observed massive proteome organization in functions related to signaling, cell adhesion, and extracellular matrix during differentiation, and less pronounced changes in processes related to innate immune response induced by the macrophage stimulation. We therefore establish our integrated differential pipeline for rapid and state-specific profiling of protein complex organization with broad utility in complex experimental designs.

## Introduction

The field of proteomics has become increasingly informative from the perspective of biology as the technology has transitioned from initially generating qualitative lists of detected proteins toward quantitative assessment of the state of the proteome over many experimental conditions in complex experimental designs^1^. However, in the cellular context functions are frequently not carried out by molecules in isolation, but rather by modules of interacting molecules^2^. A canonical example are non-covalently interacting proteins assembled into functional complexes. Large-scale protein-protein interaction (PPI) studies have demonstrated that almost all proteins participate in complexes^3^ and we have observed that the majority of the total proteome mass is assembled in stable macromolecular protein complexes^4^. The assembly state of numerous protein complexes as well as their abundance dynamically changes to respond functionally to specific environmental stimuli. To better understand the cell’s functional state, we require methods that can provide quantitative and context-dependent snapshots of the global organization of protein complexes in a way analogous to what has been achieved in more standard proteomics approaches aimed at the quantification of the expressed proteins. Methods such as affinity purification or proximity labelling combined with mass spectrometry have provided deep maps of the protein interaction space within static cellular contexts^3,5^, or alternatively, descriptions of changes for limited numbers of protein complexes in perturbed systems^6–8^. However, practical global methods aimed at monitoring complexes in many conditions have been difficult to achieve at a scale consistent with large-scale experiments needed to address complex biological questions.

Methods based on co-fractionation of native proteome extracts coupled to mass-spectrometry ^4,9– 13^ (CoFrac-MS, or protein correlation profiling - PCP) have shown substantial promise as an unbiased strategy to monitor the composition and variations of the protein complex landscape. CoFrac-MS relies on the biochemical fractionation (frequently SEC - size exclusion chromatography) of native cell protein extracts combined with identification and quantification of proteins inferred by bottom-up LC-MS/MS analysis of sequential fractions. The established data analysis concept^14–17^ rests on the idea that the identity of protein interactions, or further, the composition of protein complexes, can be inferred by reconstructing and correlating the elution patterns of individual proteins across the SEC fractionation space. Where two or more proteins co-elute, we take this as evidence of a protein interaction or complex, evidence that has to be further supported by using statistical filtering and the inclusion of orthogonal information e.g. prior knowledge that the respective proteins can interact^4^. In principle, such methods have the attractive property that they can capture a quantitative and contextual snapshot of the proteome-wide organization of proteins in modules for any given biological sample from which native protein extracts can be prepared. Comparative studies employing this analysis concept have demonstrated deep biological insights, that would have been difficult to achieve with other methods, such as the conservation of protein complex/interaction organization across metazoans^18^ and across mammalian tissues^19^, the reorganization of protein complexes as a function of cell cycle progression^16,20^, interactome disassembly during apoptosis^21^, the organization of ribosomes into polysomes^22^, and the large-scale characterization of RNA bound protein complexes^23^. While these and other studies have demonstrated the potential of the approach, the number of applied biology studies published that employ CoFrac-MS as their basis remains relatively modest compared to more standard proteomics approaches. We suggest that the explanation for the under-utilization of this apparently informative approach lies in the massive measurement resources required to complete a statistically well-powered multi-condition comparative experiment. Published studies have required weeks to months of mass spectrometer measurement time and even with such a brute force approach the number of biological conditions and experimental replicates analyzed is limited. In the course of preparing our manuscript Havugimana and colleagues proposed a method to scale such analysis using multiplex isobaric labelling^24^. A second barrier is the difficulty in extracting biologically meaningful information from the complicated high dimensional data produced by differential CoFrac-MS studies. We and others have proposed several computational strategies including an approach based on differential changes in protein SEC features between conditions (CCprofiler)^20^, an autocorrelation-based approach to detect rewiring of individual proteins across conditions (PrInCE)^15^, a PPI network centric approach that accounts for changes at multiple levels (SECAT)^16^, and a Bayesian framework to identify alterations in protein complexes (PCprophet)^17^. However, a data analysis pipeline that can simultaneously perform statistical comparisons of known protein complexes across multiple experimental conditions while also providing hypothesis free evaluation of evidence for protein complex remodeling at the individual protein level has not yet been described. We suggest that an integrated method combining increased measurement throughput with an integrated data analysis pipeline would enable the concept of CoFrac-MS to become broadly and routinely applicable in life science research.

We have recently introduced a number of advances to the CoFrac-MS approach. SEC-SWATH-MS^4,25^ employs Data Independent Acquisition (DIA/SWATH) mass spectrometry enabling reproducible, robust and sensitive quantification of peptides across protein complex fractions and experimental groups. Our analysis software CCprofiler uses prior protein connectivity information from protein complex or PPI databases to generate and execute targeted protein complex queries to detect target complexes, while controlling the error-rate using a target-decoy based statistical model^26^. This SEC-SWATH-MS strategy was used as the starting point for the developments reported in this study.

We present an integrated experimental and computational workflow for global assessment of protein complex reorganization in perturbed systems. Our approach relies on DIA/SWATH analysis of SEC fractions using short gradient chromatography that increases throughput by ~1 order of magnitude, achieving a measurement capacity of ~1 biological sample per day with similar information content compared to prior low throughput methods. The increase in throughput facilitates the comparison of multiple experimental groups with multiple biological replicates. To deal with this increase in complexity and to maximize the information content discernible from the data, we developed statistical methods to compare the data from several perspectives that we refer to as (i) assembled mass fraction – where we assess whether a given protein is shifting between monomeric and assembled states, (ii) protein-centric – where we detect and differentially quantify individual protein SEC features between conditions, and (iii) complex-centric – where we quantify changes in protein complexes detected by a hypothesis driven approach. We benchmark our workflow with respect to a typical lower throughput strategy and then demonstrate its performance by investigating rearrangements in the protein complex landscape of THP-1 human monocytic precursor cells when undergoing a phorbol ester induced differentiation into a macrophage-like phenotype^27^ and upon further induction of an inflammatory response via lipopolysaccharide (LPS) stimulation^28^ in the differentiated macrophages.

## Results

### Integrated experimental and computational workflow

To increase the throughput of the SEC-SWATH-MS workflow we optimized multiple steps of the experimental procedure (fig 1a). These included (i) parallelized sample preparation after SEC fractionation, including proteolytic digestion using 96-well FASP (Filter-Aided Sample Preparation) plates to ensure robustness and comparability, while significantly reducing sample handling steps and time^29^; (ii) direct sample loading onto solid phase extraction tips, omitting an offline reversed phase-based clean-up step; (iii) a data acquisition strategy comprising a 21-minute LC gradient (24 minute injection to injection time) using direct loading from solid phase extraction tips and embedded gradients to reduce overhead. This advance enabled the acquisition of data for 1 biological sample comprising ~60 SEC fractions per day while minimizing loss in sensitivity^30^; (iv) a DIA/SWATH acquisition strategy specifically optimized to maintain proteome coverage and quantitative robustness for short gradient analysis (Supplementary Fig. 1&2).

**Figure 1.**
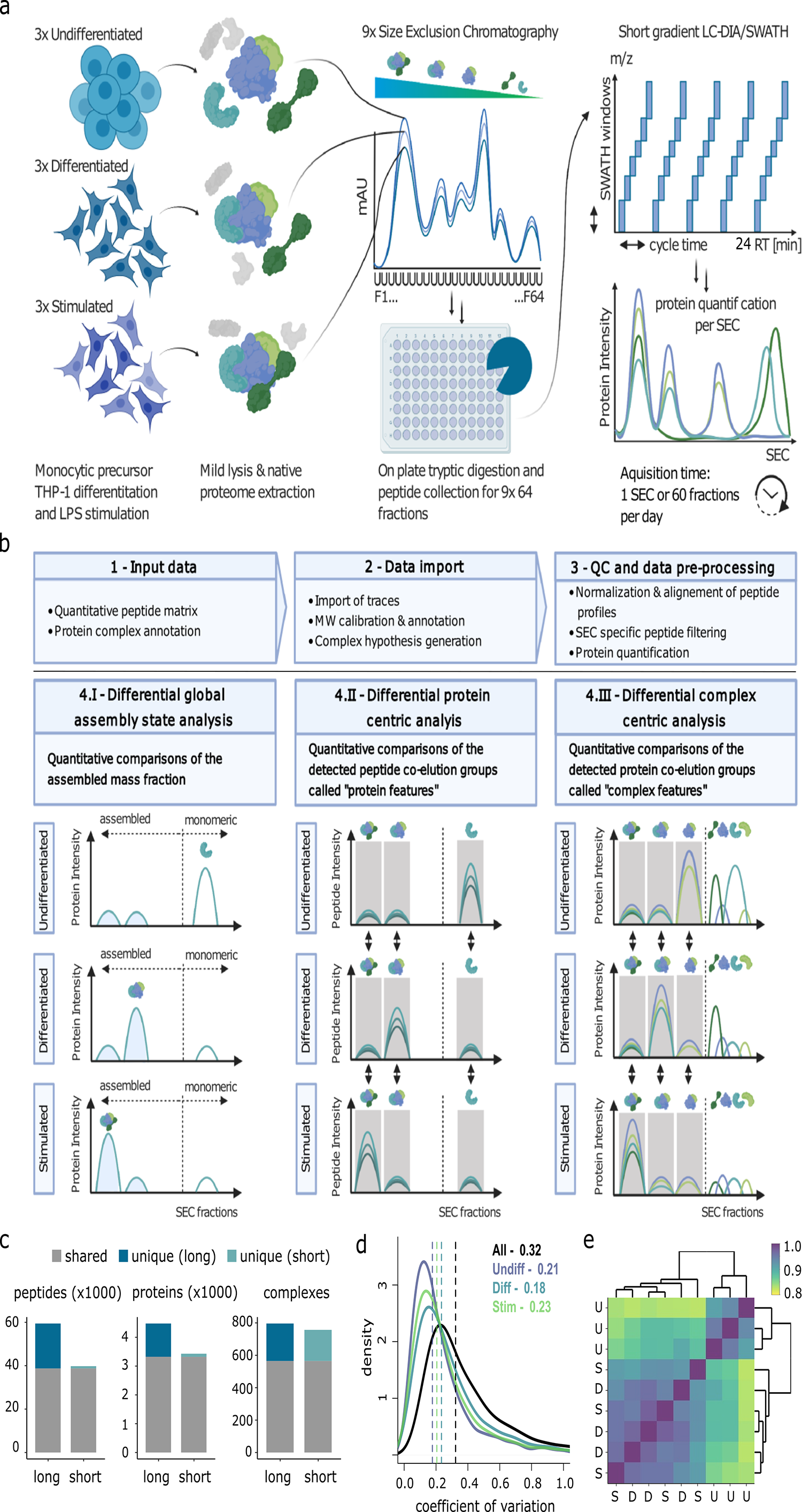
Workflow for rapid profiling of protein complex re-organization. (a) The main steps in the sample processing workflow exemplified with three biological conditions (undifferentiated, differentiated, stimulated) analyzed in triplicate. Native extracts were separated by SEC collecting 64 fractions per sample. The fractions were processed to peptides using a 96 well plate Filter Aided Sample Preparation (FASP) protocol and analyzed by 21-minutesminutes gradients in DIA/SWATH mode at a rate of 60 MS samples (~1 biological sample) measured per day. (b) The extended CCprofiler^4,20^ workflow is depicted. Steps 1,2, and 3 outline the required input data for data import that is followed by normalization, quality control procedures, and data pre-processing. Step 4 consists of quantitative comparisons between experimental groups at 3 different levels. The first differential analysis module in figure 1b panel 4.I assesses differential global assembly state analysis, reporting the relative assembled fraction compared to the monomeric state for each protein. The protein-centric analysis, in figure 1b panel 4.II reports quantitative comparisons of all detected peptide co-elution groups called protein features. Figure 1b panel 4.III depicts the CCprofiler module that supports differential complex centric analysis, where pair-wise quantitative comparisons of the detected protein co-elution groups, called complex features, between all the biological conditions are reported. (c) Benchmarking experiment using HeLa cells comparing a typical SEC-SWATH workflow using the long gradient (90 minute gradient; 126 minute injection to injection time) compared to our optimized workflow using short gradient (21 minute, 24 minute injection to injection time) analyses showing the number of peptides, proteins, or protein complexes detected. (d) CV distribution within THP-1 perturbation experimental groups compared to whole experiment. (e) Spearman correlation matrix with hierarchical clustering calculated based on SEC protein feature peak areas for undifferentiated (U), differentiated macrophages (D), and LPS stimulated (S).

In this study we benchmarked the rapid method and used it to compare THP-1 human monocytic precursor cells when undergoing a PMA-induced differentiation into a macrophage-like phenotype^27^ and subsequently a further lipopolysaccharide (LPS) stimulation of the macrophage cells to elicit an inflammatory response^28^ (Fig. 1). The native protein extracts of 3 biological replicates from all 3 biological conditions were fractionated by SEC and analyzed by DIA/SWATH mass spectrometry using short gradients as described. This amounted to 9 SEC runs of 64 fractions each, leading to a total of 576 MS runs which were acquired within 9.5 days.

Given that our optimized experimental workflow substantially increased throughput, thus facilitating the measurement of biological replicates from different experimental conditions with comparable information content, we reasoned that new algorithmic and statistical approaches were needed to fully exploit the available data and to maximize biological insight. The computational advances in the workflow are implemented in a new version of our software CCprofiler^31^ to systematically and automatically investigate changes in proteome assembly across multiple conditions or cellular states (Fig. 1b). CCprofiler includes several pre-processing functions to align SEC profiles, to compute missing values and to normalize intensities between replicates and conditions (see materials and methods). The extended CCprofiler version further enables the qualitative and quantitative detections of three complementary aspects of (differential) proteome organization, each one implemented in a specific CCProfiler module.

The first module is directed at detecting proteins that differ in their global assembly state, meaning that the relative distribution between monomeric and assembled states is different across the conditions (Fig. 1b panel 4.I). For this analysis we first exploit the log-linear relationship between the SEC elution fractions and their apparent molecular weight (Supplementary Fig. 3), enabling the assignment of a monomeric and assembled SEC elution range specific for each protein detected. The fraction of observed protein mass in the assembled SEC elution range is represented by the Assembled Mass Fraction (AMF) (see materials and methods). The differential module assesses whether a protein undergoes a significant change in AMF across the different conditions, meaning that it changes from assembled states to the monomeric state or vice versa. Importantly, AMF analysis does not require the extraction of specific elution peaks, but instead takes all fractions in the monomeric and assembled range into account, respectively. This makes the AMF analysis module also applicable to CoFrac-MS experiments of limited chromatographic resolution.

The second, protein-centric analysis module evaluates the number of distinct assembly states in which each protein is observed. We define a distinct assembly state as a resolved peptide co-elution peak group of a protein along the SEC chromatographic dimension, referred to as ‘protein feature’. Recently, we extended the protein-centric analysis to quantitatively compare protein features across different conditions^20^. In contrast to a standard differential protein expression analysis, abundance fold-changes and p-values are computed for each distinct protein feature, thereby capturing not only changes in overall protein expression, but abundance changes of specific assembly states. In addition to the feature-specific differential analysis, global differential assessment is performed by comparing integrated intensities across the entire fractionation dimension instead of restricting the analysis to a feature-specific range. The same strategies as for feature-specific estimation of log2-fold-changes and p-values are performed. Additionally, we provide the opportunity to compare the relative distribution of protein mass across the various detected assembly states (Fig. 1b panel 4.II), represented by a relative Feature-specific Mass Fraction (FMF). Here, a change in FMF across conditions indicates that the protein changes its relative distribution across different assembly states i.e. a change in the state of protein complexes that cannot be explained by a change in protein abundances only. The protein-centric differential analysis yields a fine-grained view of individual assembly states of each protein but also enables more global assessments of the overall degree of higher order assembly observed in each biological condition.

Finally, in the third analysis module CCprofiler quantitatively compares the abundances and compositions of protein complexes across different biological conditions in an automated and error-controlled manner (Fig. 1b panel 4.III). Unlike the first two strategies which are hypothesis free, the complex-centric analysis module first relies on prior protein connectivity information to query the data in a targeted fashion and to extract protein complexes based on their co-elution profiles under a controlled FDR (see materials and methods). CCprofiler then carries out a differential analysis step by comparing the signal intensity for each protein complex feature across all pairwise biological conditions. This analysis enables the consistent detection and quantitative comparison of hundreds of protein complexes across different biological conditions.

### Performance and quality assessment

To determine whether our optimized workflow had comparable information content to established CoFrac-MS strategies we benchmarked against a typical SEC-SWATH method using a 90-minute gradient (126 minute injection to injection time). In this comparison we analyzed equivalent SEC fractions from a HeLa CCL2 native protein extract with either method. The DIA/SWATH data were analyzed using Spectronaut and a previously published HeLa CCL2 spectral library^32^. We applied CCprofiler filtering and feature finding functions and evaluated the number of peptides, proteins, and protein complexes detected by both methods (Fig. 1c, Supplementary Fig. 4-5, and Supplementary Tables 1-6). Overall, we recovered 70%, 77%, and 95% of the information at the peptide, protein, and protein complex levels, respectively, when comparing the short gradient to the long gradient method with ~half order reduction in protein SEC feature dynamic range.

Having demonstrated that our rapid method still provides comparative proteome and interactome coverage, we next turned to the THP-1 perturbation experiment. We analyzed the 576 LC-MS/MS runs from the three experimental conditions in SEC triplicates using the OpenSWATH computational pipeline and a THP-1 specific spectral library generated by DDA analysis of 12 basic reversed phase fractions of a pool of the biological samples. It contained 84,453 peptide precursors, mapping to 9,375 proteins. The output, initially filtered with relaxed FDR thresholds before import to CCprofiler, contained on average 46,146 proteoytpic peptides per SEC run (range 43,785-47,297) from which we inferred 5,736 unique proteins on average (range 5,686-5,762) across the dataset at a 10% run-specific peak-group FDR, 5% global peptide FDR and 5% global protein FDR before further FDR refinement in CCprofiler (described below). We first assessed the consistency and comparability of the 9 fractionation runs by performing pair-wise alignments at the peptide-level and calculated the global correlation amongst all matching peptides (Supplementary Fig. 6). The results demonstrate that the SEC runs were reproducible and did not require further alignment. To enable a quantitative comparison of the respective SEC runs at the three analysis levels, we then normalized the intensities using a cyclic loess method (Supplementary Figure 7). To increase the confidence for downstream analyses, we filtered out peptides which were not identified in two consecutive SEC fractions, we only kept proteins supported by more than one proteotypic peptide, and we required that the remaining proteins are supported by at least two highly-correlating sibling peptides (materials and methods & Supplementary Fig. 8). After these conservative filtering steps, the mean number of detected proteins per SEC run was 4,013 (range 3,996-4,025). We next applied chromatographic feature finding to the concatenated dataset of all 9 SEC runs and detected 5,196 protein SEC elution features from 3,335 proteins at a 5% FDR threshold, amongst which 911 (27%) were detected as monomers only, whilst 2,424 (73%) had at least one elution feature in the assembled molecular weight range (> 2x monomer molecular weight). Of the 3,335 proteins 1,389 had multiple detected features at different molecular weights, thus suggesting their contributions to more than one complex (Supplementary Fig. 9-10).

To assess the global biological information content of the THP-1 dataset, we computed the coefficient of variation (CV) for peak areas of protein SEC features over the biological replicates within experimental groups and compared these to the CV over all samples across groups. Fig 1d shows the distribution of the CV for these categories where the median within experimental group CV is 0.18-0.23 whereas the median CV for all samples across the three different experimental conditions is 0.32, indicating that we have captured substantial biological variation. To further examine the global pattern of biological variability within and across experimental groups we calculated the Spearman correlation using protein SEC feature peak areas and performed hierarchical clustering (Fig 1e). This analysis showed that the undifferentiated monocytic cell state clusters distinctly from both the differentiated and stimulated states, which are not distinguished in this global analysis, indicating that the magnitude of proteome reorganization induced by differentiation is substantially higher than that of LPS stimulation of in the differentiated state.

### Protein complex re-organization in differentiated and stimulated THP-1 cells

To compare between experimental groups, we applied the 3 quantitative modules of CCprofiler beginning with the Assembled Mass Fraction (AMF) analysis. Over the 9 samples analyzed, 60-64% of the global proteome mass was estimated to be in an assembled state (Supplementary Fig. 11 and Supplementary Tables 7-8). A global view of the change in AMF with respect to the experimental comparisons is shown in Fig. 2a. Of the 3,903 proteins in the AMF analysis, 61 proteins had significant changes in their assembly state (absolute mean AMF difference larger than 0.25, BH p-value less than 0.05) comparing the differentiated to undifferentiated conditions (51 proteins increased AMF and 10 proteins decreased AMF on differentiation) and only 1 protein showed a significant change (decreased AMF) when comparing the stimulated to differentiated conditions. For example, Fig. 2b shows the average protein abundance over the SEC dimension for SHC1, a adaptor protein with broad functional roles in signaling, for each of the 3 experimental conditions. In the undifferentiated state almost all of the SHC1 signal is observed at the expected monomeric molecular weight (MMW) of ~63 kDa with only 3.7% observed at larger than 2 times the expected MMW. On differentiation we observe a clear and statistically significant shift (BH p-value = 0.003) with 32.4% of the signal for this protein observed in the assembled state. Stimulation of the differentiated cells with LPS did not produce a further significant shift in the assembly state of SHC1 with 38.4% in the assembled state (BH p-value = 0.522). SHC1 has been directly implicated as a signaling adaptor in monocyte to macrophage differentiation in previous studies^33,34^.

**Figure 2.**
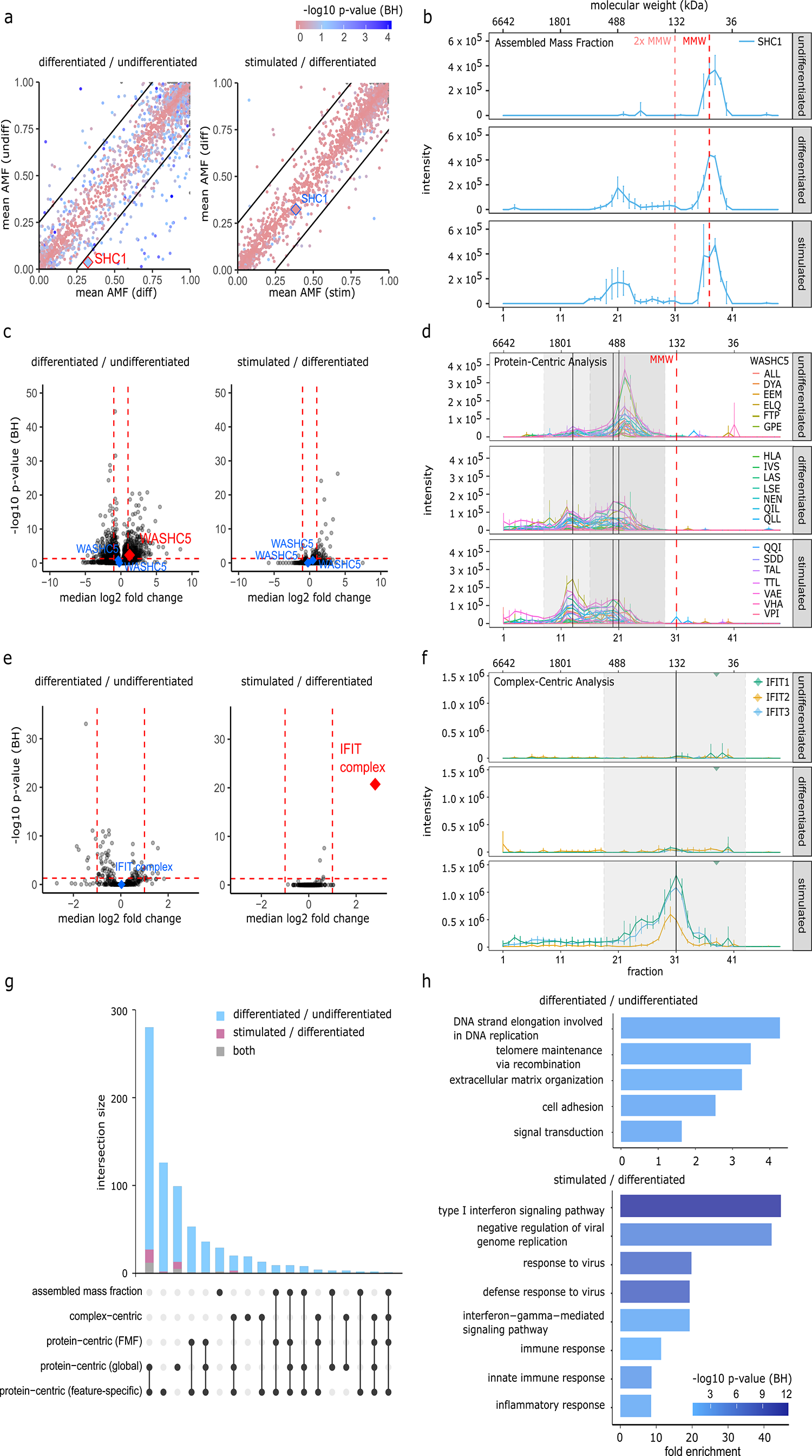
Protein complex reorganization in differentiated/stimulated THP-1 cells. (a) Scatterplot summarizing the summarizing the AMF analysis in the pairwise comparisons of interest. Points on the diagonal indicate no change between conditions and points outside the thresholds indicate an effect size > 25% with the BH p-value indicated by the pink/blue color scale. SHC1 is highlighted with a significantly altered AMF in differentiated versus undifferentiated cells (red) but not in stimulated versus differentiated cells (blue). (b) Absolute mass fraction (AMF) analysis showing protein SEC profiles for SHC1 in the 3 experimental conditions, demonstrating an AMF shift from the monomeric state toward an assembled state, going from undifferentiated monocytes to differentiated macrophages. MMW and 2x MMW indicate the expected monomeric molecular weight and twice the expected monomeric molecular weight. Biological replicates are collapsed to median ± SD. (c) Volcano plot summarizing the protein-centric analysis where each data point represents one protein SEC feature. Highlighted points are WASHC5 protein SEC features that do (red) or do not (blue) pass significance thresholds. (d) Protein-centric analysis showing Peptide SEC traces for WASHC5 in the 3 experimental conditions demonstrating a reduction in abundance of the lower molecular weight SEC feature and an increase in the higher molecular weight SEC feature going from the undifferentiated condition toward the differentiated condition. Grey background indicates SEC feature boundaries and black lines indicate feature apex. Biological replicates are collapsed to mean ± SD. (e) Volcano plot summarizing the complex-centric analysis where each data point represents one detected protein complex. Highlighted points are the IFIT complex that changes significant in the stimulated vs. differentiated comparison (red) but not in the differentiated vs. undifferentiated comparison (blue). (f) Complex-centric analysis showing protein SEC profiles for the 3 components of the IFIT complex where the complex is observed in the stimulated condition but not in the undifferentiated or differentiated conditions. Grey background indicates SEC feature boundaries and black lines indicate feature apex. (g) Upset plot showing the overlap in proteins deemed to have significant changes in each of the quantitative comparisons in either the differentiated vs. undifferentiated comparison (blue), the stimulated vs. differentiated comparison (pink), or shared in both (grey) (h) Functional enrichment analysis both comparisons of interest based on combined set of proteins significant from all comparison methods.

We next applied the Protein-Centric Analysis module of CCprofiler to the THP-1 dataset. Overall, we observed feature-specific quantitative differences (absolute log2FC larger than 1, BH p-value less than 0.05) between SEC features for 540 proteins in the differentiated vs. undifferentiated comparison and in 30 proteins for the stimulated vs. differentiated comparison (Fig 2c and Supplementary Tables 9-11). In the protein-centric global comparison, where the signal from all SEC fractions for a given protein is summed and compared across conditions, we observed significant changes for 428 proteins in the differentiated vs. undifferentiated comparison and 39 proteins for the stimulated vs. differentiated comparison. We then performed the feature-specific mass fraction (FMF) comparison in which we can infer whether a protein changes its relative distribution across different assembly states. We detected 114 proteins in the differentiated vs. undifferentiated comparison that underwent changes in their assembly state that were not attributable to changes in overall protein quantity and, in contrast, we could detect no proteins in this category for the stimulated vs. differentiated comparison. Fig. 2d shows SEC profiles for peptides mapping to WASHC5, a component of the WASH complex associated with endosome regulation. We observe 2 distinct protein SEC features both substantially in excess of the expected MMW, indicating the likely participation of WASHC5 in two distinct complexes. A significant reduction of peak area of the lower molecular weight feature in the undifferentiated vs. differentiated conditions is observed in combination with an apparent (although not significant) increase in the higher molecular weight feature. In Fig. 2c the summary for all protein features is shown in volcano plots for the comparisons of interest with the protein features for WASHC5 highlighted. While the specific role of the WASH complex in differentiation is not broadly understood mouse cells lacking the WASH complex were shown to be deficient in haemopoietic differentiation including at the transition from monocyte to macrophage lineages^35^ and therefore reorganization of this complex is plausibly functionally relevant in this monocyte to macrophage transition.

Finally, we applied the complex-centric module of CCprofiler to compare the set of detected protein complexes between conditions. To generate a comprehensive input hypothesis-set for the complex-centric analysis, we merged the CORUM database^36^ with the String database^37^ partitioned to create discrete protein complex hypotheses using the ClusterONE algorithm^38^, originally created to detect potentially overlapping protein complexes from PPI datasets (Supplementary Table 12). This resulted in 3,127 complex hypotheses, from which 644 were detected; 104 were fully detected, 375 were detected with at least 50% of the subunits present and 165 were identified with less than 50% of the subunits present, all with a 5% FDR at the complex-detection level (Supplementary Fig. 12). We further collapsed these 644 confidently detected protein complex queries to 321 likely unique protein complexes based on subunit composition and position in the SEC dimension^25^. Overall, we observed significant quantitative changes in 17 protein complexes, composed of 73 protein subunits, in the differentiated vs. undifferentiated comparison. This is in contrast to the stimulated vs. differentiated comparison in which only a single protein complex was called as significantly different (Fig. 2e and Supplementary Tables 13-17). Fig. 2f shows the protein level SEC traces for the 3 annotated subunits of the IFIT complex. The IFIT complex, composed of the Interferon-induced protein with tetratricopeptide repeats 1-3, is a well described factor in interferon-induced response with particular functional relevance in anti-viral function although more recent data implicates the IFIT complex in regulatory function in inflammatory reponses^39^. For example, LPS stimulation in macrophages has been shown to increase IFIT expression levels in order to enhance the secretion of proinflammatory cytokines including TNF alpha and IL-6^40^. We observe a clear signal for the IFIT complex at the expected molecular weight in the LPS stimulated condition and compared with only baseline amounts in the differentiated and undifferentiated conditions. As such, the increased expression of the IFIT proteins, and their assembly into a complex, is consistent with the LPS stimulation employed in our experiment. Interestingly, IFIT1 and IFIT3 (but not IFIT2) also appear in an as yet unannotated higher molecular weight assembly.

Fig. 2g summarizes the number and overlap of proteins that we call as significantly changing in each of the CCprofiler analysis modes described for both experimental comparisons (Supplementary Table 18). As expected, there is some redundancy between these modes of analysis but also much complementarity as each strategy is tuned to detect different aspects of protein complex re-organization. This view of the data underscores the magnitude difference in the response from the perspective of proteome organization to the chosen biological perturbations. Here we expect to capture changes related both to increases in the abundance of protein complexes driven primarily by changes in the abundance of their protein subunits (likely exclusive to ‘protein-centric (global)’, ‘protein-centric (feature-specific)’, and ‘complex-centric’ categories), as well as changes in protein assembly composition (exclusive to ‘protein-centric (FMF)’ and ‘assembled mass fraction’, or combinations of those mechanisms that appear in multiple categories). This graph also underscores the observation that substantially more changes occur in cells that transition from the suspension monocyte state to the adherent macrophage state than in the comparison of macrophage cells that are stimulated to upregulate immune defenses by LPS stimulation. To obtain a global functional picture of the response to these perturbations from the perspective of proteome organization we performed a functional enrichment analysis based on the consolidated list of proteins significantly altered in each element of the CCprofiler analysis. Fig. 2h shows the results for both comparisons where we find enriched terms that are consistent with differentiated vs undifferentiated comparison (i.e. extracellular matrix organization, cell adhesion, signal transduction, etc.) and the stimulated vs. differentiated comparison (i.e. type I interferon signaling pathway, innate immune response, etc.). We examined the distribution of protein SEC feature peak areas and determined that while the peak areas of protein features that are detected at >2x the expected monomer molecular weight follow the same peak area distribution as all detected protein SEC features, the set of protein SEC features that match to protein complexes detected in the complex-centric analysis are shifted toward higher peak areas indicating we are somewhat less likely to successfully call complexes for lower abundance features (Supplementary Fig. 13). This observation underscores the utility of assessing the data in a protein complex hypothesis free manner, as in the AMF and protein-centric analysis modes, in addition to the complex centric analysis.

In order to render the data easily accessible and viewed in depth, and enable manual query of community-based testing of novel putative interacting proteins supported by the presence of coelution profiles, we have made the data accessible to the community via a web portal^32^ (https://collins-lab.shinyapps.io/secexplorer_thp1/). This online tool provides the opportunity to manually query the SEC profiles of our 3 biological THP-1 conditions by providing an interactive viewing. It enables the manual query for locally co-eluting proteins to potentially identify *de novo* interactions and to visualize the results from the 3 differential CCprofiler modules.

## Discussion

Methods using co-fractionation as a basis have promised characterization of the state of protein complexes and their reorganization upon cellular perturbation in a global and quantitative fashion. However, with the current state of the workflows this remains a somewhat distant goal with respect to routine application, especially for complex experimental designs. While a number of studies employing this approach have demonstrated substantial biological insight, the general strategy has failed to break into mainstream use. At the outset of this study we identified two major factors holding back progress, namely, (i) the resources required per biological sample for typical implementations of this strategy are not practically compatible with complex study designs including multiple experimental conditions with biological replication, and, (ii) a lack of integrated software solutions that could perform differential statistical analysis at all levels of interest (assembled mass fraction, protein-centric, and complex-centric). In this study we address both barriers by developing an integrated experimental and computational pipeline for rapid quantitative profiling of protein complex states.

We present an optimized method that facilitates the interrogation of many biological samples in perturbation experiments with biological replicates within a feasible time-frame and a robust data analysis pipeline. By using robust short gradient liquid chromatographic separation and DIA/SWATH data acquisition we could reduce the MS acquisition time of 9 SEC runs of 64 fractions each, totaling 576 samples down to 9.6 days. This is in comparison to an estimated 50.4 days for the same project based on the 90 minute gradient and 126 minute injection to injection time that we used in the long gradient comparison for this study, although we note many studies using this strategy perform 2-4 hour gradients for CoFrac-MS analyses^13^. In real terms the actual increase in throughput is substantially higher because the short gradient chromatography using solid phase extraction tips for loading and embedded gradients for separation reduces substantially the need for maintenance procedures such as instrument cleaning and column changes that typically interrupt data acquisition blocks using classical long gradient methods. As such, this represents an order of magnitude reduction in the time required to acquire data for this type of experiment and does not require the complexity and experimental design constraints associated with multiplex labelling. While the gains in throughput are critical to the further development of this strategy, we expect commensurate benefits to data quality as the inevitable effect of drift in instrument performance over time will be substantially mitigated by the reduced measurement time and reduced need for instrument maintenance during data acquisition^30^. In a benchmarking experiment using HeLa cells we demonstrate that the information content from our rapid method is comparable to that of a standard long gradient approach, and we demonstrate in our THP-1 perturbation experiment that the across-group variation in our quantitative data exceeds the within-group variation indicating we capture biological information. Further, since our data was acquired a number of improvements in MS data acquisition schemes aimed at maximizing the numbers of peptides/proteins quantified in short gradient data have been introduced and we expect our strategy to directly benefit from these, increasing sensitivity and protein complex coverage and reducing analysis time^41–43^.

We made several algorithmic improvements, embedded in several novel modules of our CCprofiler^4,20^ software pipeline, that maximize the information extracted from the more complex experimental designs that are facilitated by our higher throughput method. These include the capability to assess between group differences from 3 perspectives that each have their own advantages/disadvantages. Complex-centric analysis provides the richest information on the reorganization of protein complexes as a function of the perturbation but is likely missing many interesting changes because it relies on the prior information in the form of testable protein complex hypotheses that may be incomplete. We note that several tools have been introduced recently that leverage machine learning to define protein complex hypotheses from CoFrac-MS data^14,15,17^ and these could be used as input for the CCprofiler complex-centric analysis. The protein-centric strategy is free of any such assumptions and simply asks whether a given protein feature in the SEC dimension changes between experimental groups. Our results show that this comparison can sensitively detect changes that are a proxy for changes in protein complex re-organization or abundance in a fine-grained manner. The assembled mass fraction strategy similarly does not require background protein complex information and further does not require feature finding in the SEC dimension, meaning that it may detect changes in proteins/complexes which smear across many SEC fractions that would be missed by the other methods.

The biological perturbations that we chose induce different cellular states that likely rely on quite different molecular mechanisms and this difference is reflected in our results. We observed evidence for a significantly higher number of changes in protein organization or abundance when comparing the monocyte state to the differentiated macrophage state than when comparing the unstimulated versus stimulated macrophage cells. This is also apparent from the unsupervised clustering which clearly shows the undifferentiated state as clearly distinguished from the differentiated and LPS stimulated states. Differentiation from a suspension monocyte-like phenotype to an adherent macrophage-like state is a gross and irreversible phenotypic change that likely requires the remodeling of many protein complexes that are visible to our method. When integrating changes observed at the three levels of CCprofiler analysis we see substantial alterations in proteome organization relating to broadly to signaling, cell adhesion, and extracellular matrix related functions. Whereas the induction of an inflammatory response may be better characterized as a change in signaling/activation in given pathways that rely more on PTMs (post translational modifications) or transient changes in protein complex assembly state that are more difficult to detect. This observation underscores the idea that data generated from this approach would benefit from combination with other data types (e.g. proteoforms or PTMs) where their interdependence could be assessed^44,45^. Nevertheless, while a smaller number of changes were observed in macrophage LPS stimulation as compared with differentiation from monocyte to macrophage, functional categories related to innate immune response were clearly overrepresented in the results integrated from our three level CCprofiler analysis.

With the introduction of our rapid integrated method, we anticipate that global profiling of protein complex re-organization in perturbation experiments with complex experimental designs will be enabled as a primary tool in systems biology research and beyond.

## Data and software availability

The mass spectrometry proteomics data have been deposited to the ProteomeXchange Consortium via the PRIDE^46^ partner repository with the dataset identifier PXD036711.

Reviewer access pre-publication is available via https://www.ebi.ac.uk/pride/ Username

Password: AGZSowSQ

The CCprofiler software including differential analysis modules is available at https://github.com/CCprofiler/CCprofiler/tree/differential. Analysis scripts for this paper are at https://github.com/ibludau/THP_SEC_SWATH_MS. Code for the SECexplorer instance to view the THP1 data is implemented via R Shiny and is available at https://github.com/collins-ben/SECexplorer_THP1.

## Author contributions

I.B., C.N., C.M., R.A., and B.C.C. conceptualized and designed the study; C.N., C.M., and P.X. performed experiments; I.B. conceived the data analysis software; I.B., C.N., C.M., and B.C.C performed the data analysis; M.H. built the web portal and made conceptual contributions; A.F., F.U., and F.F. devised the plate-based sample preparation method; R.A. and B.C.C supervised the study; I.B., C.N., C.M., and B.C.C wrote the manuscript with contributions from all authors.

## Acknowledgements

B.C.C. was supported by an Ambizione grant (PZ00P3_161435) of the Swiss National Science Foundation (SNSF), and by Queen’s University Belfast establishment grant. C.M. was supported by grant #2017-402 of the Strategic Focal Area “Personalized Health and Related Technologies (PHRT)” of the ETH Domain. I.B. was supported by a Swiss National Science Foundation Postdoc.Mobility fellowship (P400PB_191046). R.A. lab was funded by the ERC (ERC 20140AdG 670821).

## Materials and methods

### Cell culture

The human monocytic cell line THP-1 (LGC, ATCC-TIB-202) was cultured and expanded in RPMI 1640 media (Gibco, 61870-010) supplemented with 10% FCS (BioConcept, 2-01F00-I) and 1% Penicillin/Streptomycin (Gibco, 15140-122) and kept at a confluency between 0.5 - 1.2 × 10^6^ cells per ml at 37°C in a 5% CO_2_ incubator. 1.5 × 10^6^ THP-1 cells were differentiated when supplemented with 50 ng/mL PMA (Sigma, P1585) for 48h and when stated, the differentiation treatment included a 24-hour stimulation with 100 ng/mL LPS (Sigma, L2630). The suspension cells or differentiated adherent cells were washed with PBS (Gibco, 10010-023) and were sedimented in a pellet by centrifugation at 300g kept at 4°C. The cell pellets were immediately snap-frozen in liquid nitrogen.

### Sample preparation for library generation

The proteins were extracted from the frozen cell pellets by lysing the cells with 1% SDC (Sigma, D6750) in HNN Buffe pH 7.8 (50 mM HEPES, 150 mM NaCl, 50 mM NaF, 200 μM Na_3_VO_4_, 1 mM PMSF, 1x Protease Inhibitors (Sigma, P8215), 1x Benzonase (Sigma, E1014)), and incubated for 5 minutes at room temperature. The lysates were centrifugated at 13’000g for 10 minutes to remove insoluble materials. The extracted proteins were reduced at 5mM TCEP for 30 minutes at 37°C while shaking at 500 rpm and subsequently alkylated in 10 mM Iodoacetamide for 30 minutes at 37°C. The proteins were precipitated overnight in 100% Acetone at -20°C and pelleted by a 30-minute centrifugation step at 4°C. The protein pellets were then resuspended in 1% SDC, 8M Urea in 0.1 M Ammonium bicarbonate and sonicated for 10 minutes. The proteins were diluted to 0.1 M ammonium bicarbonate and digested overnight with Trypsin (Promega, V5113) at 37°C with a protein-to-enzyme ratio of 50:1. The digestions were stopped with 50% TFA and the SDC was removed by two centrifugation steps of 10 minutes each at 16’000g. The peptides were desalted and cleaned-up using C18 columns (The Nest Group, #SEM SS18V) and were resuspended in 5% acetonitrile, 0.1% formic acid with iRT peptides (Biognosys, Ki-3002).

For the spectral library generation, a fraction of all samples was pooled together, dried using a vacuum centrifugation at 45°C and resuspended in Buffer A (20 mM ammonium formate, 0.1% ammonia solution, pH 10). 200 μg of peptides were injected into an Agilent Infinity 1260 (HP Degasser, Vial Sampler, Cap Pump) and 1290 (Thermostat, FC-μS) system and separated on a 25 cm long C18 reverse-phase column (YMC Triart) with 3μm particle size and 12 nm of pore size. The peptides were separated at a flow rate of 12μl/min by a linear 56-min gradient from 5% to 35% Buffer B (20 mM ammonium formate, 0.1% ammonia solution, 90% acetonitrile in water, pH 10) against Buffer A (20 mM ammonium formate, 0.1% ammonia solution, pH 10) followed by a linear 4-min gradient from 35% to 90% Buffer B against Buffer A and 6 min at 90% Buffer B. The resulting 36 fractions were pooled into 12 samples. The buffer of the pooled samples was evaporated using vacuum centrifugation at 45 °C and the resulting 12 samples were resuspended in 2% ACN, 0.1% FA with iRT peptides (Biognosys).

### SEC protein complex extraction and fractionation

Protein complexes fractionation was performed as previously described ^31^. THP-1 cells were thawed and lysed in mild conditions by homogenization with a lysis buffer composed by 0.5% NP-40 detergent and protease and phosphatase inhibitors (50 mM HEPES pH 7.5, 150 mM NaCl, 0.5% NP-40, 1 mM PMSF, 400 nM Na3CO4, protease inhibitors cocktail (Sigma-Aldrich, MI, USA)). Cell debris and membranes were removed by 15 minutes of ultracentrifugation (55,000×g, 4 °C) and the detergent was removed by 30 kDa molecular weight cut-off membrane and exchanged with the SEC buffer (50 mM HEPES pH 7.5, 150 mM NaCl). The samples were concentrated for a final protein concentration between 7-12 μg/μl. After 5 min of centrifugation at 16,900 ×g at 4 °C, the supernatant was directly injected to a Yarra-SEC-4000 column (300 × 7.8 mm, pore size 500 Å, particle size 3 μm, Phenomenex, CA, USA). 0.8 mg of native proteome extract (estimated by Pierce™ BCA Protein Assay Kit, Thermo Fisher Scientific, MA, USA) was injected for each SEC run at 4 °C with a flow rate of 500 μl/min, for a total chromatographic time of 30 min. Fraction collection was performed in the retention time window from 10 to 26 min, at 0.25 min per fraction, for a total of 64 fractions collected.

The molecular weight calibration curve for SEC fractionation was obtained by running a protein standard mix (Column Performance Check Standard, Aqueous SEC 1, AL0-3042, Phenomenex, CA, USA) before each sample injection (Supplementary Table 19).

### Sample preparation for Mass Spectrometry analysis

Sample processing for bottom-up analysis of SEC fractions was performed on a 96-well plate MWCO filters (AcroPep Advance Filter Plates for Ultrafiltration 1mL Omega 10K MWCO; Pall Corporation, USA) ^47^. Prior to usage, the filters are washed twice with 200 μl of water that was successively removed by centrifugation at 1800 g for 30 min. 64 fractions for each sample (total fraction volume 125 μl) were loaded and concentrated on the filters through centrifugation, until the complete removal of the SEC buffer.

Protein denaturation and reduction was obtained incubating the samples at 37 °C for 30 min with 5 mM of TCEP in 8M Urea/20 mM ammonium bicarbonate (AMBIC) (pH 8.8). Alkylation of cysteine residues was performed adding a final concentration of 50 mM IAA/20 mM AMBIC and incubating in the dark and room temperature for 1 h. After the reaction, the plates were centrifuged for removing the Urea buffer and washed for three times with 20 mM AMBIC. Protein digestion was carried out at 37 °C for 16 h, adding to each well 1 μg of trypsin (Promega, Switzerland) and 0.3 μg of Lysyl Endopeptidase (Mass Spectrometry grade, FUJIFILM Wako Pure Chemical Industries, Japan). The resulting peptides were collected by centrifugation and the plates were washed once more with 100 ul of ddH20.

### LC-MS analysis

DIA/SWATH analysis of the peptide fractions was performed on Evosep One system (Evosep Biosystems, Denmark) ^30^ coupled to an AB Sciex TripleTOF 6600 instrument (Sciex, MA, USA) equipped with a NanoSpray III ion source (Sciex). The samples are loaded in Evotips (Evosep Biosystems, Denmark), after resuspension in solvent A (0.1% FA water solution, Fisher Scientific AG, Switzerland) and the addition of iRTs peptides (Biognosys) in a ratio 1:100 for the retention time alignment requested for SWATH acquisition. 75% of the peptide recovered from each SEC fraction was loaded. For the loading, the C18 stage tips (Evotips) were soaked with 100 μl of 2-propanol during the activation and the conditioning steps. The activation step consisted in the washing with 20 μl of solvent B (0.1 % FA in ACN, Fisher Scientific AG, Switzerland), followed by the conditioning with 20 μl of solvent A. Prior the sample loading step, 10 μl of solvent A is added on top of the tips, ensuring that the tips remain wet during the loading step. For each steps, the Evotips were centrifuged for 1 min at a speed of 700 g for the elution of the solvents. The last step (i. e. washing step) was performed using 100 μl of solvent A, and the loaded tips are added with 200 μl of solvent A for preserving the samples during the entire injection of the batch.

The separation of peptides was performed selecting the “60 samples per day” method, consisting in 24 minutes of total cycle time, for 21 minutes of gradient length, 3 minutes of overhead time at a flow rate of 1 μl/min. A partial gradient is applied (0-35% solvent B) in order to elute the peptides from the Evotip by two couples of low pressure pumps. The peptides were then pushed in a C-18 nanoConnect LC column (8 cm column, ID 100 μm packed with 3 μm Reprosil, PepSep, Denmark) using an high pressure pump and solvent A ^30^. The ESI coupling was obtained using a Nano Source Emitter Stainless Steel Nano-bore 1/32 (Thermo Fisher Scientific).

The ESI tuning parameters were the following: spray voltage, 2800 V; ion source gas flow (GS1), 16; curtain gas flow (CUR), 35; interface heater temperature (IHT), 100°C and declustering potential, 100.

The Evosep system was controlled by the Axel Semrau Chronos software (Axel Semrau GmbH, Germany), while the mass spectrometer acquisition software was Analyst TF 1.7.1 (Sciex).

Data-independent acquisition (SWATH/DIA) mass spectrometry^48^ was performed for the quantitative analysis of the 576 SEC fractions (64 fractions per sample) obtained from the 9 SEC experiment. SWATH scan performed using an updated scheme of 64 variably sized precursor co-isolation windows ^49^, covering similar precursor densities (in terms of number and intensity) within all SWATH windows. The SWATH windows cover the precursors ions in the range of 350-1500 m/z and 350-1500 in the MS2 SWATH scans, the accumulation time was 100 ms for the MS1 and 20 ms for each SWATH window, resulting in a cycle time of 1.38 s. For fragmentation, it was applied a rolling collisional energy with a collisional energy spread of 15 eV.

### DDA MS analysis for the library generation

The 12 high pH fractioned peptide samples were separated on an Eksigent nanoLC Ultra AS2 1D Plus and expert 400 autosampler system (Eksigent, Dublin, CA) coupled to a TripleTOF 5600 through a NanoSpray III ion source using a Data Dependent Acquisition (DDA) scheme. The 20 cm long nanoLC column was packed in house using a 75 μm inner diameter PicoFrit emitter (New Objective, Woburn) with Magic C18 AQ 3 um, 200 Å particles. The separation was performed at room temperature with a flow rate of 300 nl/min. All the LC solvents were all of mass spectrometry grade. The LC solvent A was composed of 98% water, 2% acetonitrile and 0.1% formic acid, LC solvent B was 98% acetonitrile, 2% water and 0.1% formic acid. The peptides were eluted over 120 minutes, with a linear gradient from 5% to 35% LC solvent B. One MS1 scan with a m/z range of 360-1460 and an accumulation time of 250 ms was followed by 20 MS2 scans with m/z ranges of 50-2000 and accumulation times of 100 ms. The dynamic exclusion time was set to 20 seconds.

### Data processing

#### DDA data analysis for the library generation

DDA-MS data acquired from peptide fractionation of the full THP-1 cell lysates (see above) were processed for the SWATH library generation following the protocol previously described ^50^.

MS spectra were searched for peptide matches against the human UniProt/SwissProt reference database (reviewed, canonical entries, June 2017) using Comet 2018.01 rev. 0 MS/MS search engine. The search was carried out using trypsin cleavage, 30 ppm precursor and 0.05 Da fragment ion mass tolerance, carbamidomethyl (C) as static and oxidation (M) as variable modification and a maximum of 2 enzyme missed cleavages. The results from the search were statistically scored using Peptide Prophet (statistical validation of PSMs) and iProphet (peptide sequence validation) of the Trans-Proteomic Pipeline (TPP v5.0.0 POLAR VORTEX rev 0), filtering the results at 1% peptide FDR (0.815939 iprob) as determined using the tool Mayu ^51^. A wider peptide-level FDR cut-off (5% FDR on protein level, compared to requiring 1% FDR) was chosen in order to increase sensitivity for the recovery of true positive peptide signals.

The resulting spectra were then gathered for the generation of the consensus spectra library using SpectraST including retention time calibration. The 6 most abundant fragment ion transitions per precursor from the b_n_ or y_n_ ion series were selected, with a m/z range of 350-2000 and aa fragment charge states 1-2. The final library contains query parameters for 506,717 precursors of 73,007 peptides mapping to 9375 protein groups. Moreover, to the spectra consensus library reverse decoy (506,581 decoys transitions) were generated for the FDR scoring provided by the SWATH/DIA data analysis workflow.

### DIA/SWATH data analysis

For the THP-1 experiment the DIA/SWATH data collected from the analysis of SEC fractions were analyzed through peptide-centric analysis, querying 506,717 fragment precursors from the sample-specific peptide library generated (see above) in the SWATH MS2 spectra, using OpenSWATH v2.1^52,53^ PyProphet and TRIC^54^ workflow. Initially, one global classifier was trained on a subsampled set of SEC fractions across the experiment using pyProphet-cli^55^. Peptides from all fractions were then quantified and scored using the pre-trained scoring function using pyProphet and TRIC. The HeLa benchmark dataset was analysed with Spectronaut v14 using a previously published HeLa CCL2 spectral library^32^.

### CCProfiler

The first differential analysis module in CCprofiler is tailored towards detecting proteins that differ in their global assembly state, meaning that the relative distribution between monomeric and assembled protein mass is different across the conditions. Since this module depends on the assignment of the fractionation dimension into a monomeric and assembled range based on the monomeric molecular weight of each protein, the analysis is currently only available for SEC datasets and requires both a molecular weight calibration of the fractions and a monomeric molecular weight annotation of the measured proteins. The cutoff between the monomeric and assembled SEC range is set at the fraction corresponding to two times the expected monomeric molecular weight of a protein. Based on this initial division of the SEC dimension, the assembled mass fraction (AMF) of each protein can be estimated by the fraction of the detected MS signal in the assembled mass range relative to the total, globally detected signal:

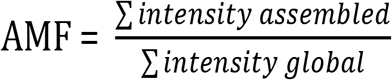

A change in AMF is subsequently estimated by the difference in mean AMF across conditions:

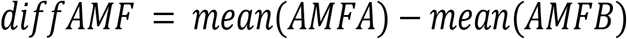

Here, AMFA and AMFB denote the AMF values of two conditions A and B. Since AMF values are not normally distributed and bound by zero and one, a conventional t-test for significance estimation is not applicable. Instead, CCprofiler applies a beta-regression model and p-value estimation by a likelihood-ratio test to derive significance estimates (for details see below). Multiple testing correction is performed by Benjamini-Hochberg adjustment of the derived p-values ^56^. Proteins with significant adjusted p-values and large AMF differences, are indicated to have a different proportion of individual proteins associated to higher order assemblies across the conditions. Notably, this information is derived independent from any feature (i.e. peak group) detection and does not require knowledge of the protein’s exact interaction partners.

### Differential analysis of distinct protein assembly states and detection of protein rewiring

To further gain insights into distinct protein assembly states, we have previously introduced the protein-centric analysis concept for CoFrac-MS data within a single condition ^31^. Here, we extend the protein-centric analysis concept to enable the differential assessment of distinct protein assembly states. To achieve consistent protein feature (i.e. peptide co-elution peak group) detection across conditions and replicates, peptide-level traces are first integrated by summing the intensities across all samples in the provided tracesList. The integrated traces are subsequently used for protein-centric feature finding, applying random peptide assignments as decoy model for p- and q-value estimation ^31^. Each protein can thereby be assigned to potentially multiple distinct assembly states, as indicated by the detection of multiple unique protein features. Following this initial protein feature detection, differential analysis is performed to compare the signal intensity within each protein feature across conditions.

Differential analysis is performed in 5 steps: (i) Peptide-level intensities are computed for each protein feature and sample. Missing values in single fractions, replicates or conditions are imputed by uniformly sampling values between zero and the minimum detected signal of a peptide. The peptide intensity of one feature is then calculated by summing the intensities of all fractions across the corresponding protein feature range. (ii) The mean intensity across all replicates within a condition (specified by the design matrix) is calculated. (iii) The log2-fold-change between conditions is calculated based on the mean feature intensities. (iv) If replicates are available, p-values are estimated by comparing the summed intensities across conditions by a non-paired t-test. If no replicates are available, p-values are estimated by comparing each fraction within a feature by a paired t-test across the conditions. (v) To subsequently derive protein-level information, the peptide-level tests are aggregated as follows: (1) protein log2-fold-changes are derived from the median log-2-fold change across all detected peptides of the protein, and (2) protein p-values are estimated by determining the fold-change adjusted median p-value and applying a beta distribution as described by Teo et al. ^57^ and Suomi et al. ^58^ (for details see method section). (vi) Multiple testing correction is performed by Benjamini-Hochberg adjustment of the protein-level p-values ^31^.

In addition to the feature-specific differential analysis, global differential assessment is performed by comparing integrated intensities across the entire fractionation dimension instead of restricting the analysis to a feature-specific range. The same strategies as for feature-specific estimation of log2-fold-changes and p-values are performed. To assess whether the signal within a protein feature is changing because of a global change in the protein’s expression or due to a rearrangement of the proteins relative distribution across different assembly states, an additional analysis step is available in CCprofiler. Here, the relative feature-specific mass fraction (FMF) is estimated by the fraction of the detected MS signal in the feature-specific mass range relative to the total detected signal:

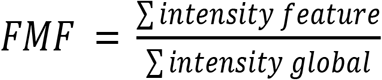

A change in FMF is subsequently estimated by the difference in mean FMF across conditions:

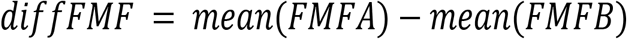

Here, FMFA and FMFB denote the FMF values of two conditions A and B. Similar to the concept introduced for comparing AMF values, CCprofiler applies a beta-regression model and p-value estimation by a likelihood ratio test ^59^ to derive significance estimates for the change in FMF across conditions (for details see methods section). Since the initial assessment of FMF values is performed on peptide-level data, protein-level information is derived by aggregation across all detected peptides as follows: (1) FMF differences are derived from the median diffFMF across all detected peptides of the protein, and (2) p-values are estimated by determining the difference adjusted median p-value and applying a beta distribution as described by Teo et al. ^57^ and Suomi et al. ^58^(for details see method section). Multiple testing correction is performed by a Benjamini-Hochberg adjustment of the p-values ^56^. A significant change in the FMF across conditions indicates that the protein’s relative contribution to different distinct assembly states has changed across the conditions, thus providing insights into protein rewiring which is not observable by global proteome analyses. In contrast to complex-centric analyses, described in the following section, protein-centric differential analysis enables the assessment of changes in distinct protein assembly states independent of actually knowing the protein’s exact interaction partners.

### Protein complex detection and differential analysis

The final analysis module in CCprofiler is focused on the complex-centric detection and differential assessment of protein complexes. We have previously introduced the basic concept of complex-centric analysis for CoFrac-MS data of a single condition ^31^. In summary, prior protein connectivity information is used to query CoFrac-MS data directly for evidence of pre-defined complexes. By using random protein assignments as a decoy model for error rate estimation, complex-centric analysis enables the detection of hundreds of protein complexes at high sensitivity and under controlled FDR. Here, we expand the complex-centric analysis strategy to allow the quantitative comparison between complexes detected across different cellular conditions. Analogous to the protein-centric workflow described in the previous section, protein-level traces are first integrated by summing the intensities across all samples in the provided tracesList to ensure consistent signal detection across conditions and replicates. The integrated traces are subsequently used for complex-centric feature detection. Only the most complete complex feature (i.e. protein co-elution peak group) for each complex query is considered for scoring and FDR estimation. After filtering for q-values (e.g. 0.05), the complex features are appended by secondary features with high correlation values (peak correlation 0.7). These secondary features can for example entail potential sub-complexes or complex variants ^31^.

Following this initial protein complex feature detection, a differential analysis step can be performed to compare the signal intensity within each complex feature across different conditions. The analysis concept is analogous to the differential analysis strategy implemented on the level of protein features (see previous section). The initial differential testing is performed on peptide level, while results are subsequently aggregated on the protein level. For complex-centric analysis, the protein-level results are additionally aggregated to the complex level, again following the same strategy as compared to aggregation from peptide to protein level. Finally, multiple testing correction is performed by a Benjamini-Hochberg adjustment of the p-values ^56^.

### P-value estimation for AMF and FMF differences

P-value estimation for AMF and FMF differences was performed by first transforming the AMF and FMF (y) to values between zero and one, while excluding the extremes (0 and 1) ^60,61^:

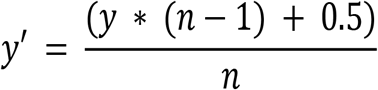

Here, n denotes the sample size, which was six for the presented dataset. The resulting y’ values were used for fitting a beta-regression model with the betareg R package with default parameters ^60,61^. The lrtest function of the lmtest R package ^59^ was subsequently used for p-value estimation by a likelihood-ratio test with default parameters. Multiple testing correction was performed by the p.adjust function of the stats base package, using the “fdr” method corresponding to correction by Benjamini-Hochberg ^56^.

### P-value estimation for aggregating peptide-level tests to the protein and complex level

Peptide-level p-values were aggregated to the protein-level by applying the strategy presented by Teo at al. ^57^ and Suomi et al. ^58^. First the median of peptide-level p-values is used as a score for each protein taking the direction of change into account. The protein-level significance of the detection is subsequently calculated using a beta distribution ^58^. The same strategy is applied to aggregate protein-level p-values to the complex level. Multiple testing correction is performed by the p.adjust function of the stats base package, using the “fdr” method corresponding to correction by Benjamini-Hochberg ^56^.

### CCprofiler analysis workflow and parameters

All R-scripts for the CCprofiler analysis are openly available on github. The following provides a summary of the most important processing steps and the selected parameters for the presented analysis.

Due to the very low molecular weight of later SEC factions, the data was limited do fractions 1 to 49 for CCprofiler analysis. Missing peptide intensity values (for which both the previous and following fraction contained measured intensity values) were imputed by a spline fit across the SEC dimension. After missing value imputation, peptide intensity values were normalized across conditions and replicates by applying a cyclic loess normalization^16,62,63^. Low-confidence peptides were subsequently removed, keeping only peptides with (1) at least three consecutive detections across any replicate, (2) at least one high correlating sibling peptide (maximum correlation >= 0.5), and (3) a good average sibling peptide correlation (>= 0.2). Protein quantification was performed by summing the top two most intense peptides consistently across all replicates.

To determine proteins with a significant change in their assembly state across conditions, a mean difference in AMF of >= 30% and a Benjamini-Hochberg adjusted p-value <= 0.05 were required.

Protein-centric analysis was performed with following parameters: corr_cutoff=0.9, window_size=7, rt_height=1, smoothing_length=7, perturb_cutoff=“5%” and collapse_method=“apex_only”. Only protein features passing the 5% FDR threshold were further considered. For the differential analysis, a minimum log2 fold-change of one and a Benjamini-Hochberg corrected p-value of 0.05 were required for significance in all pairwise analyses. To determine protein features with a significant change in their relative abundance in comparison to the total protein intensity across conditions, a mean difference in FMF of >= 30% and a Benjamini-Hochberg adjusted p-value <= 0.05 were required.

For complex-centric analysis, we first defined a set of target protein complex queries. This was achieved by combining queries derived from CORUM^36^ and StringDB^37^. We derived protein complex queries from StringDB v10 (9606.protein.links.v10.txt). Protein identifiers were mapped to Uniprot accessions via BioMart. The interactions were filtered for a minimal combined_score of 980. We applied the ClusterONE algorithm ^38^ for PPI network partitioning with following parameters: d=0.95. Weights were set to the combined_score divided by 1000. CORUM derived protein complex queries were taken directly from within the CCprofiler package ^31^. The complex queries were combined and decoys were generated randomly by requiring a minimum edge distance of 3. Complex-centric analysis was performed with following parameters: corr_cutoff=0.9, window_size=7, rt_height=1, smoothing_length=7, perturb_cutoff=“5%” and collapse_method=“apex_network”. Only complex features with a molecular weight higher than two times the largest monomeric molecular weight of any of its participating subunits were considered. For each protein complex query, the complex feature with the highest number of participating subunits was selected for FDR estimation, filtering for a maximum FDR of 5%. Secondary features were appended to the final results based on a minimum peak correlation threshold of 0.7. To reduce redundancy across the detected complex features between different queries, features were collapsed with following parameters: rt_height = 0 and distance_cutoff = 1.25. For the differential analysis, a minimum log2 fold-change of one and a Benjamini-Hochberg corrected p-value of 0.05 were required for significance in all pairwise analyses.

