## supplementary text for "Rapid profiling of protein complex re-organization in perturbed systems"

### Supplementary Figures

A

| Condition | Column length (cm) | Flow rate (nl/min) | Gradient lenght (min) | N variable windows | MS² scan time (msec) | MS² mass range | Sample load (µgs) |
|---|---|---|---|---|---|---|---|
| A) Standard DIA/SWATH method | 20 | 300 | 90 | 64 | 50 | 50-2000 | 1 |
| B) Shorten column | 10 | 300 | 90 | 64 | 50 | 50-2000 | 1 |
| C) Increase flowrate | 10 | 1000 | 90 | 64 | 50 | 50-2000 | 1 |
| D) Short gradient-Method 1 | 10 | 1000 | 20 | 64 | 25 | 50-2000 | 1 |
| E) Short gradient-Method 2 | 10 | 1000 | 20 | 32 | 50 | 50-2000 | 1 |
| F) Short gradient-Method 3 | 10 | 1000 | 20 | 100 | 15 | 50-2000 | 1 |
| G) Short gradient-Method 4 | 10 | 1000 | 20 | 64 | 25 | 350-1500 | 1 |
| H) Short gradient-Method 5 | 10 | 1000 | 20 | 64 | 25 | 50-2000 | 2 |

B

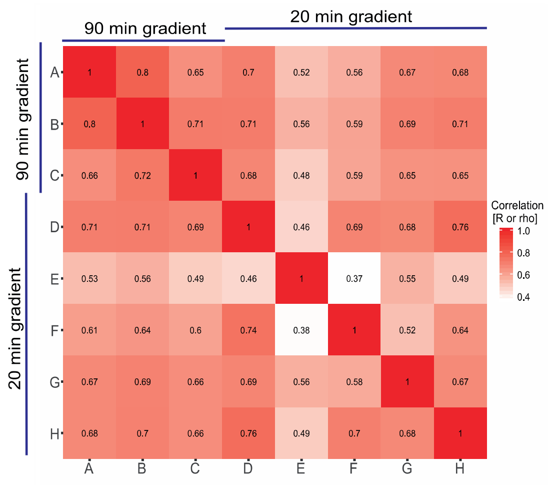


C

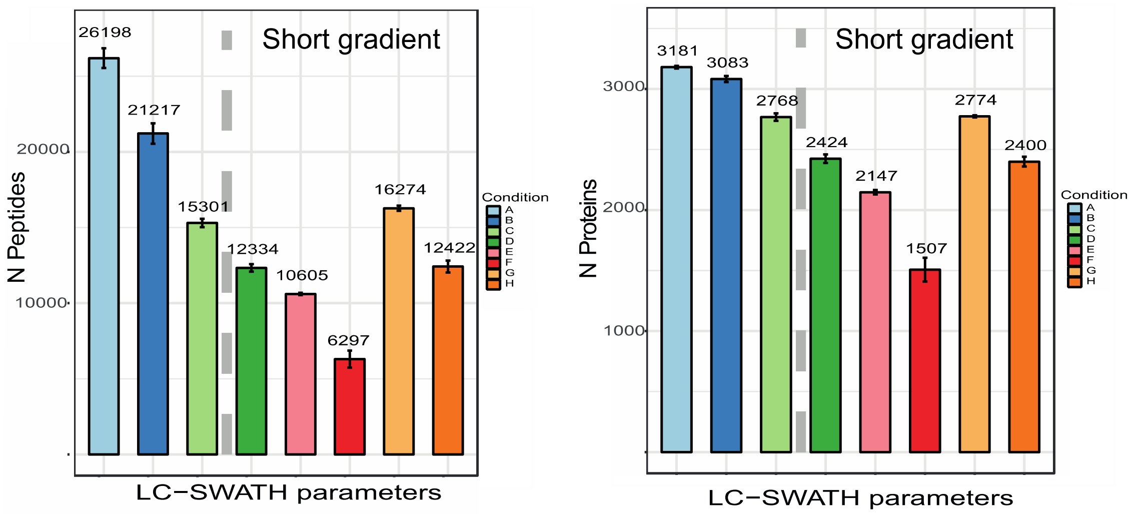


**Supplementary Figure 1 – Effect of chromatographic optimization on peptide detection rates**

Effect of the column length, flow rate, gradient length and number of DIA/SWATH variable windows in the DIA/SWATH performances (sample correlation, number of peptides and proteins identification). All the

runs were acquired with Eksigent nanoLC Ultra AS2 1D Plus and expert 400 autosampler system (Eksigent, Dublin, CA) coupled to a TripleTOF 5600 (Sciex) through a NanoSpray III ion source (Sciex). The 20 cm and 10 cm nanoLC columns were packed in house using a 75 µm inner diameter PicoFrit emitter (New Objective, Woburn) and ProntoSIL 3 um, 200 Å particles (Bischoff Chromatography, Germany).
For the chromatographic separation an aqueous solution with 2% ACN/0.1 FA % was used as buffer A, while a 98 % ACN%/0.1% FA solution as buffer B.

**Panel A. Short scheme of the different conditions tested.**

A) Standard DIA/SWATH method using a scheme of 64 variably sized precursor co-isolation windows. The SWATH windows cover the precursor ions in the range of 360-1460 m/z and 50-2000 in the MS2 SWATH scans, the accumulation time was 250 ms for the MS1 and 50 ms for each SWATH window, resulting in a cycle time of 3.5 s. For fragmentation, it was applied a rolling collisional energy with a collisional energy spread of 15 eV. 1 µg of peptides were loaded in the sample loop. The separation was carried out with a 20-cm capillary column at 300 nl/min flow for a 90-min gradient from 5 to 30% of buffer B.

B) Shorten column using a scheme of 64 variably sized precursor co-isolation windows. The SWATH windows cover the precursors ions in the range of 360-1460 m/z and 50-2000 in the MS2 SWATH scans, the accumulation time was 250 ms for the MS1 and 50 ms for each SWATH window, resulting in a cycle time of 3.5 s. For fragmentation, it was applied a rolling collisional energy with a collisional energy spread of 15 eV. 1 µg of peptides were loaded in the sample loop. The separation was carried out with a 10-cm capillary column at 300 nl/min flow for a 90-min gradient from 5 to 30% of buffer B.

C) Increased separation flowrate using a scheme of 64 variably sized precursor co-isolation windows. The SWATH windows cover the precursors ions in the range of 360-1460 m/z and 50-2000 in the MS2 SWATH scans, the accumulation time was 250 ms for the MS1 and 50 ms for each SWATH window, resulting in a cycle time of 3.5 s. For fragmentation, it was applied a rolling collisional energy with a collisional energy spread of 15 eV. 1 µg of peptides were loaded in the sample loop. The separation was carried out with a 10-cm capillary column at 1000 nl/min flow for a 90-min gradient from 5 to 30% of buffer B.

D) Short gradient-Method 1. 64 variably sized precursor co-isolation windows scheme. The SWATH windows cover the precursors ions in the range of 360-1460 m/z and 50-2000 in the MS2 SWATH scans, the accumulation time was 250 ms for the MS1 and 25 ms for each SWATH window, resulting in a cycle time of 1.9 s. For fragmentation, it was applied a rolling collisional energy with a collisional energy spread of 15 eV. 1 µg of peptides were loaded in the sample loop. The separation was carried out with a 10-cm capillary column at 1000 nl/min flow for a 20-min gradient from 5 to 30% of buffer B.

E) Short gradient-Method 2. 32 variably sized precursor co-isolation windows scheme. The SWATH windows cover the precursors ions in the range of 360-1460 m/z and 50-2000 in the MS2 SWATH scans, the accumulation time was 250 ms for the MS1 and 50 ms for each SWATH window, resulting in a cycle time of 1.9 s. For fragmentation, it was applied a rolling collisional energy with a collisional energy spread of 15 eV. 1 µg of peptides were loaded in the sample loop. The separation was carried out with a 10-cm capillary column at 1000 nl/min flow for a 20-min gradient from 5 to 30% of buffer B.

F) Short gradient-Method 3. 100 variably sized precursor co-isolation windows scheme. The SWATH windows cover the precursors ions in the range of 360-1460 m/z and 50-2000 in the MS2 SWATH scans, the accumulation time was 250 ms for the MS1 and 15 ms for each SWATH window, resulting in a cycle time of 1.8 s. For fragmentation, it was applied a rolling collisional energy with a collisional energy spread of 15 eV. 1 µg of peptides were loaded in the sample loop. The separation was carried out with a 10-cm capillary column at 1000 nl/min flow for a 20-min gradient from 5 to 30% of buffer B.

G) Short gradient-Method 4. 64 variably sized precursor co-isolation windows scheme. The SWATH windows cover the precursors ions in the range of 360-1460 m/z and 350-1500 in the MS2 SWATH scans, the accumulation time was 250 ms for the MS1 and 25 ms for each SWATH window, resulting in a cycle time of 1.9 s. For fragmentation, it was applied a rolling collisional energy with a collisional energy spread of 15 eV. 1 µg of peptides were loaded in the sample loop. The separation was carried out with a 10-cm capillary column at 1000 nl/min flow for a 20-min gradient from 5 to 30% of buffer B.

F) Short gradient-Method 5. 64 variably sized precursor co-isolation windows scheme. The SWATH windows cover the precursors ions in the range of 360-1460 m/z and 50-2000 in the MS2 SWATH scans, the accumulation time was 250 ms for the MS1 and 50 ms for each SWATH window, resulting in a cycle time of 3.5 s. For fragmentation, it was applied a rolling collisional energy with a collisional energy spread of 15 eV. 2 µg of peptides were loaded in the sample loop. The separation was carried out with a 10-cm capillary column at 1000 nl/min flow for a 90-min gradient from 5 to 30% of buffer B.

**Panel B. Pearson (upper triangle) and Spearman correlation (lower triangle) of fragment ions intensities across the tested conditions (90 min gradient vs. 20 min gradient).**

Effect of the column length, flow rate, gradient length and number of DIA/SWATH variable windows in terms of the correlation of the total fragment intensities. The SWATH-MS data were analyzed using OpenSWATH, PyProphet and TRIC workflow (see Methods section). No further normalization was applied. From the plot results clear as the SWATH acquisition method [i.e. SWATH number of windows; Condition E (32 SWATH windows) vs Condition D (64 SWATH windows) and Condition F (100 SWATH windows)] influences most the performances of the analysis with respect that the column length (i.e. Condition A vs Condition B) and the flow rate (i.e. Conditions A/B vs Condition C).

**Panel C. Number of peptides (left panel) and number of proteins (right panel) identified across the tested conditions.**

Effect of the column length, flow rate, gradient length and number of DIA/SWATH variable windows in terms of total number of peptides and proteins identified.

The short gradient decreases the number of peptides identified by 53% (Condition A vs Condition D), whereas of 24% the total proteins compared to the long gradient. As shown for the correlation analysis, the major effect observed for short gradients is determined by the selection of the number of SWATH windows and the MS2 scan range. The 100 SWATH window method (Condition F) showed 76% and 53% of peptides and protein loss, respectively, with respect to the standard conditions (Condition A). The smaller MS2 scan range (Condition H) improved the performances of 24% and 13% of peptides and proteins characterized, respectively, in comparison with the broader MS2 scan range (Condition D).

A

| Condition | N variable windows | MS² scan time (msec) | MS² mass range | Cycle time (msec) |
|---|---|---|---|---|
| **A** | 32 | 40 | 350-1500 | 1430 |
| **B** | 32 | 40 | 50-1500 | 1430 |
| **C** | 32 | 40 | 50-2000 | 1430 |
| **D** | 64 | 20 | 350-1500 | 1430 |
| **E** | 64 | 20 | 50-1500 | 1430 |
| **F** | 64 | 20 | 50-2000 | 1430 |

B

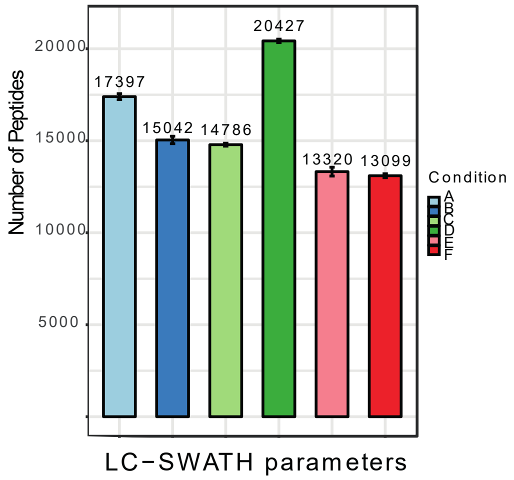


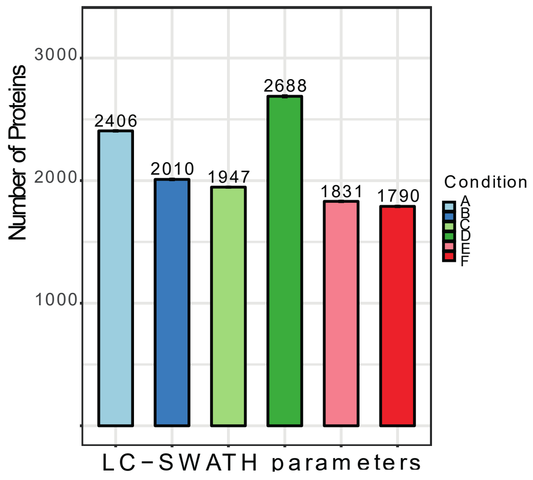


C

| Condition | N variable windows | MS² scan time (msec) | MS² mass range | Cycle time (msec) |
|---|---|---|---|---|
| **A** | 16 | 80 | 350-1500 | 1430 |
| **B** | 24 | 54 | 350-1500 | 1446 |
| **C** | 32 | 40 | 350-1500 | 1430 |
| **D** | 40 | 32 | 350-1500 | 1430 |
| **E** | 48 | 27 | 350-1500 | 1446 |
| **F** | 56 | 23 | 350-1500 | 1438 |
| **G** | 64 | 20 | 350-1500 | 1430 |

D

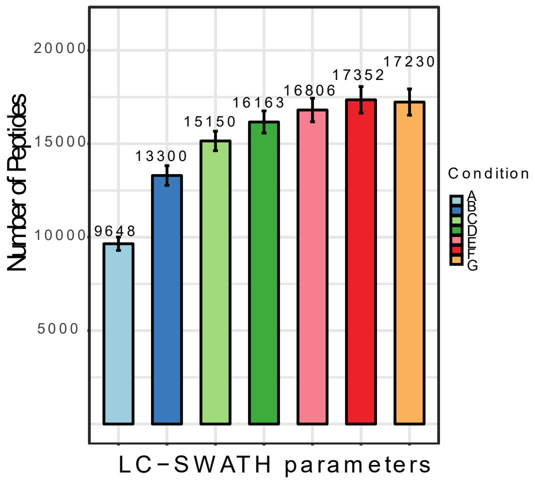


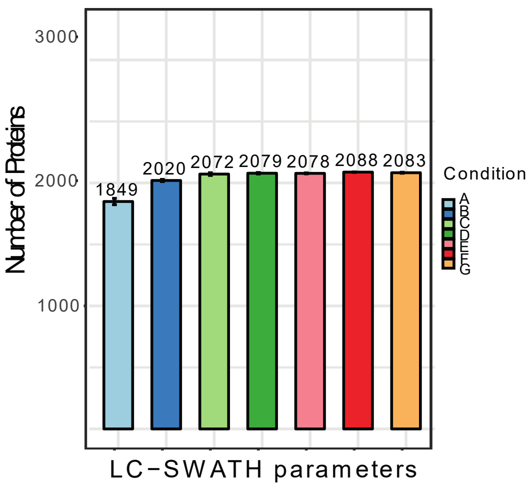

**Supplementary Figure 2 - Effect of DIA/SWATH optimization on peptide detection rates**

Effect of the MS2 scan range (Panels A and B) and number of SWATH windows (Panels C and D) in the DIA/SWATH performances (number of peptides and proteins identified). All the runs were performed on Evosep One system couple to an TripleTOF 5,600 instrument equipped with a NanoSpray III ion source selecting the "60 samples per day" method and using the nanoConnect LC column (8 cm column, ID 100 µm packed with 3 µm Reprosil, PepSep) (see Methods section).

Panel A. Short scheme of the different conditions tested for the MS2 scan range.

A) 32 variably sized precursor co-isolation windows. The SWATH windows cover the precursors ions in the range of 350-1460 m/z and 350-1500 in the MS2 SWATH scans, the accumulation time was 100 ms for the MS1 and 40 ms for each SWATH window, resulting in a cycle time of 1.430 s. For fragmentation, it was applied a rolling collisional energy with a collisional energy spread of 15 eV. 1 µg of peptides were loaded in the sample loop.

B) 32 variably sized precursor co-isolation windows with the MS2 SWATH scans range of 50-1500. The other parameters were the same of Condition A.

C) 32 variably sized precursor co-isolation windows with the MS2 SWATH scans range of 50-2000. The other parameters were the same of Condition A.

D) 64 variably sized precursor co-isolation windows. The SWATH windows cover the precursor ions in the range of 350-1460 m/z and 350-1500 in the MS2 SWATH scans, the accumulation time was 100 ms for the MS1 and 20 ms for each SWATH window, resulting in a cycle time of 1.430 s. For fragmentation, it was applied a rolling collisional energy with a collisional energy spread of 15 eV. 1 µg of peptides were loaded in the sample loop.

E) 64 variably sized precursor co-isolation windows with the MS2 SWATH scans range of 50-1500. The other parameters were the same of Condition D.

F) 64 variably sized precursor co-isolation windows with the MS2 SWATH scans range of 50-2000. The other parameters were the same of Condition D.

Panel B. Number of peptides (left panel) and number of proteins (right panel) identified across the conditions tested for the MS2 scan range.

Effect of the MS2 scan range in the DIA/SWATH performances in terms of number of peptides and proteins identified.

The smaller MS2 scan range improves the numbers of peptides and proteins identified both with 32 and 64 SWATH windows methods. For the 32 SWATH windows method, the 350-1500 MS2 scan range (Condition A) improves of ~14% and ~16% the number of peptides and proteins identifications (with respect of conditions B and C). For the 64 SWATH windows method, the 350-1500 MS2 scan range (ConditionD) improves of ~35% and ~32% the number of peptides and proteins identifications (with respect of conditions E and F). As shown here, the influence of the MS2 scan range in the mass spectrometry acquisition increases with an increased number of SWATH windows; a smaller mass range improves the sensitivity of the detection and allowing a higher number of events for the same full cycle time (1.430 secs).

Panel C. Short scheme of the different conditions tested for the number of SWATH windows.

A) 16 variably sized precursor co-isolation windows. The SWATH windows cover the precursor ions in the range of 350-1460 m/z and 350-1500 in the MS2 SWATH scans, the accumulation time was 100 ms for the MS1 and 80 ms for each SWATH window, resulting in a cycle time of 1.430 s. For fragmentation, it was applied a rolling collisional energy with a collisional energy spread of 15 eV. 1 µg of peptides were loaded in the sample loop.

B) 24 variably sized precursor co-isolation windows. The accumulation time for each SWATH window is 54 ms, for a total cycle time of 1.446 s. The other parameters were the same of Condition A.

C) 32 variably sized precursor co-isolation windows. The accumulation time for each SWATH window is 40 ms, for a total cycle time of 1.430 s. The other parameters were the same of Condition A.

D) 40 variably sized precursor co-isolation windows. The accumulation time for each SWATH window is 32 ms, for a total cycle time of 1.430 s. The other parameters were the same of Condition A.

E) 48 variably sized precursor co-isolation windows. The accumulation time for each SWATH window is 27 ms, for a total cycle time of 1.446 s. The other parameters were the same of Condition A.

F) 56 variably sized precursor co-isolation windows. The accumulation time for each SWATH window is 23 ms, for a total cycle time of 1.438 s. The other parameters were the same of Condition A.
G) 64 variably sized precursor co-isolation windows. The accumulation time for each SWATH window is 20 ms, for a total cycle time of 1.436 s. The other parameters were the same of Condition A.

Panel D. Number of peptides (left panel) and number of proteins (right panel) identified across the conditions tested for the number of SWATH windows. Evaluation of the performances of different DIA/SWATH methods in function of the number of variable SWATH windows. The shorter gradient
The smaller MS2 scan range improves the numbers of peptides and proteins identified both with 32 and 64 SWATH windows methods. For the 32 SWATH windows method, the 350-1500 MS2 scan range (Condition A) improves of ~14% and ~16% the number of peptides and proteins identifications (with respect of conditions B and C). For the 64 SWATH windows method, the 350-1500 MS2 scan range (ConditionD) improves of ~35% and ~32% the number of peptides and proteins identifications (with respect of conditions E and F). As shown here, the influence of the MS2 scan range in the mass spectrometry acquisition increases with an increased number of SWATH windows; a smaller mass range improves the sensitivity of the detection and allowing a higher number of events for the same full cycle time (1.430 secs).

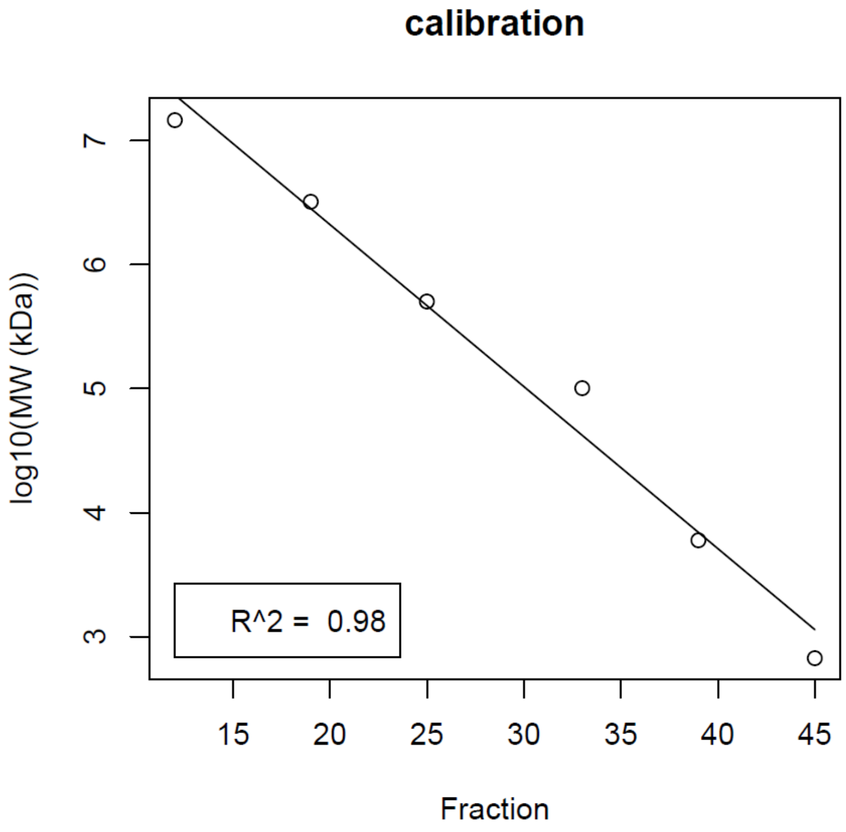


**Supplementary figure 3 – SEC molecular weight calibration**

Example of a molecular weight calibration based on the log-linear relationship of the calibrant's molecular weights and the elution fraction in the SEC separation.

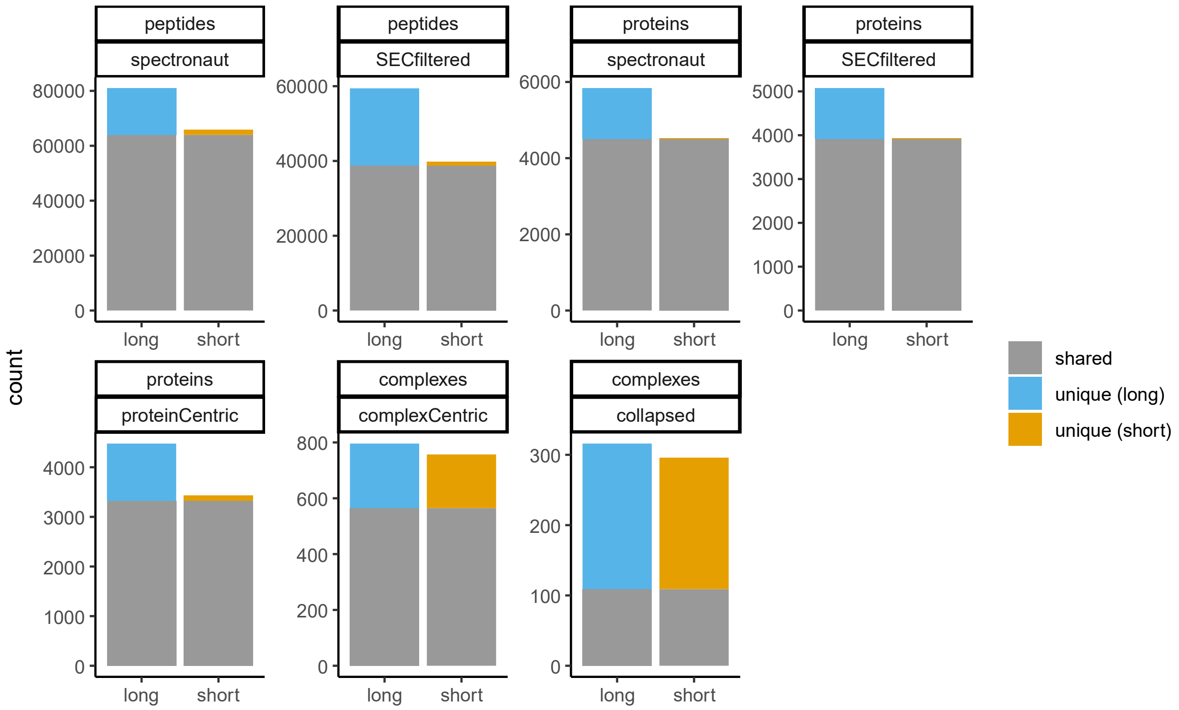


**Supplementary figure 4 – Benchmarking of rapid method in HeLa cells**

Comparison of our newly developed rapid method using short gradient chromatography ('short') versus a representative standard SEC-SWATH method ('long'). The number of peptides, inferred proteins, or inferred complexes detected (shared or uniquely in each method) using either the long or short method are shown. 'Spectronaut' refers to detections at the stated thresholds after Spectronaut analysis and 'SECFiltered' refers to more stringent filtering utilizing consecutive fraction and at least 2 correlated sibling peptides. 'Collapsed' refers to protein complexes after removing redundancy due to multiple complexes hypotheses detecting the same/similar complex features. See methods for further description of categories depicted.

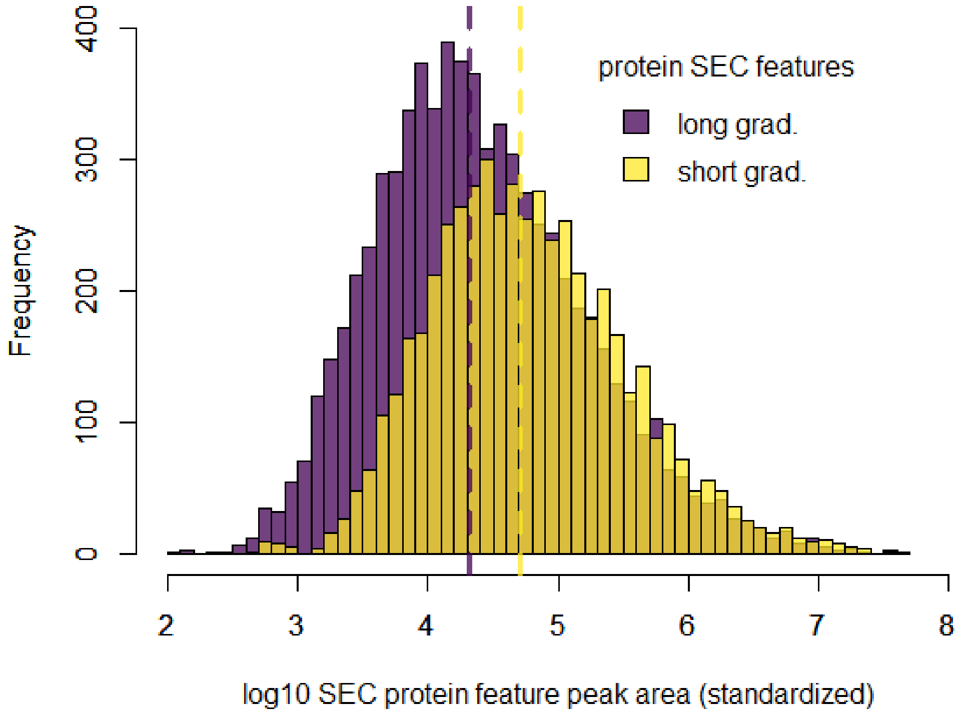


**Supplementary figure 5 – Dynamic range assessment in short versus long gradient**
Distribution of protein SEC features in the short or long gradient analysis HeLa Benchmark.

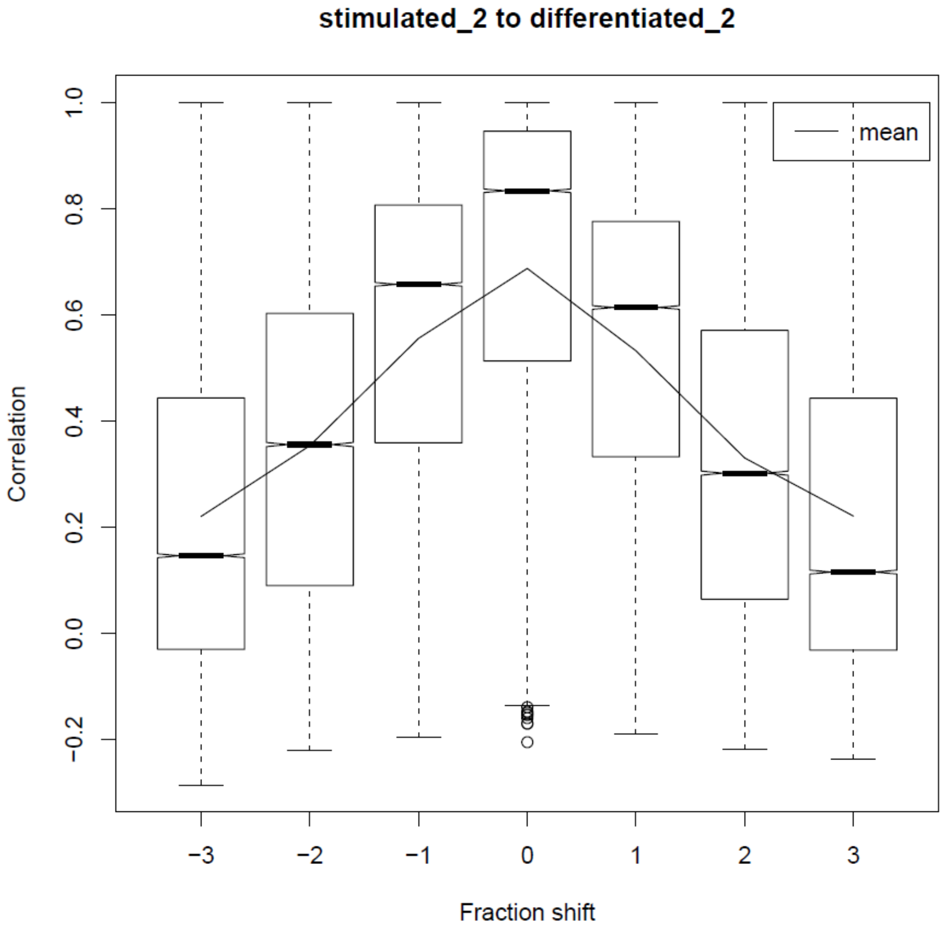


**Supplementary figure 6 – Pairwise SEC alignment**
Representative example of a pair-wise alignment between the peptide-level SEC traces to illustrate different fraction shift offsets.

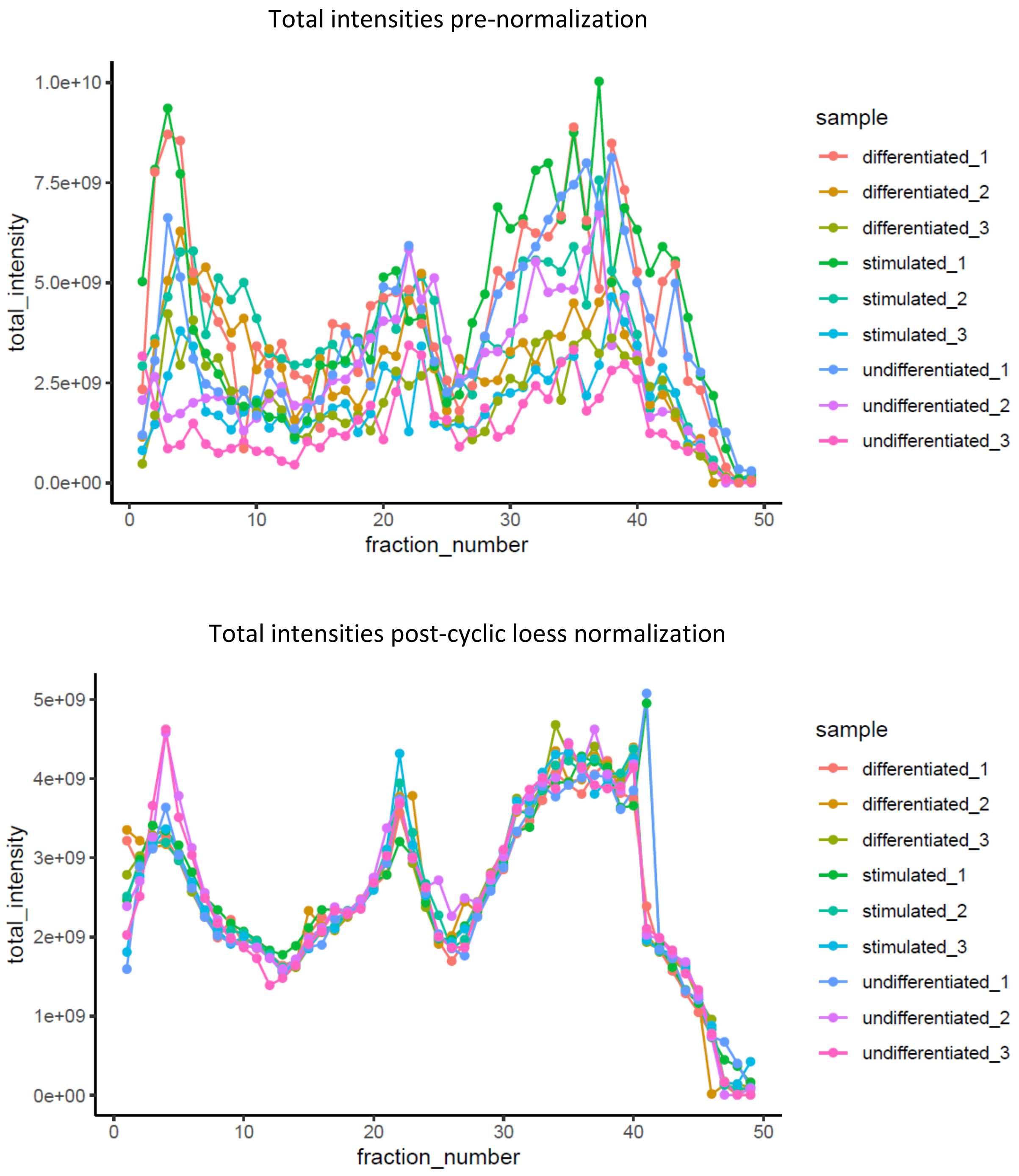


**Supplementary figure 7 – Cyclic loess normalization**

Collective total ion intensities for all 9 SEC runs before and after cyclic loess normalization for all 9 runs.

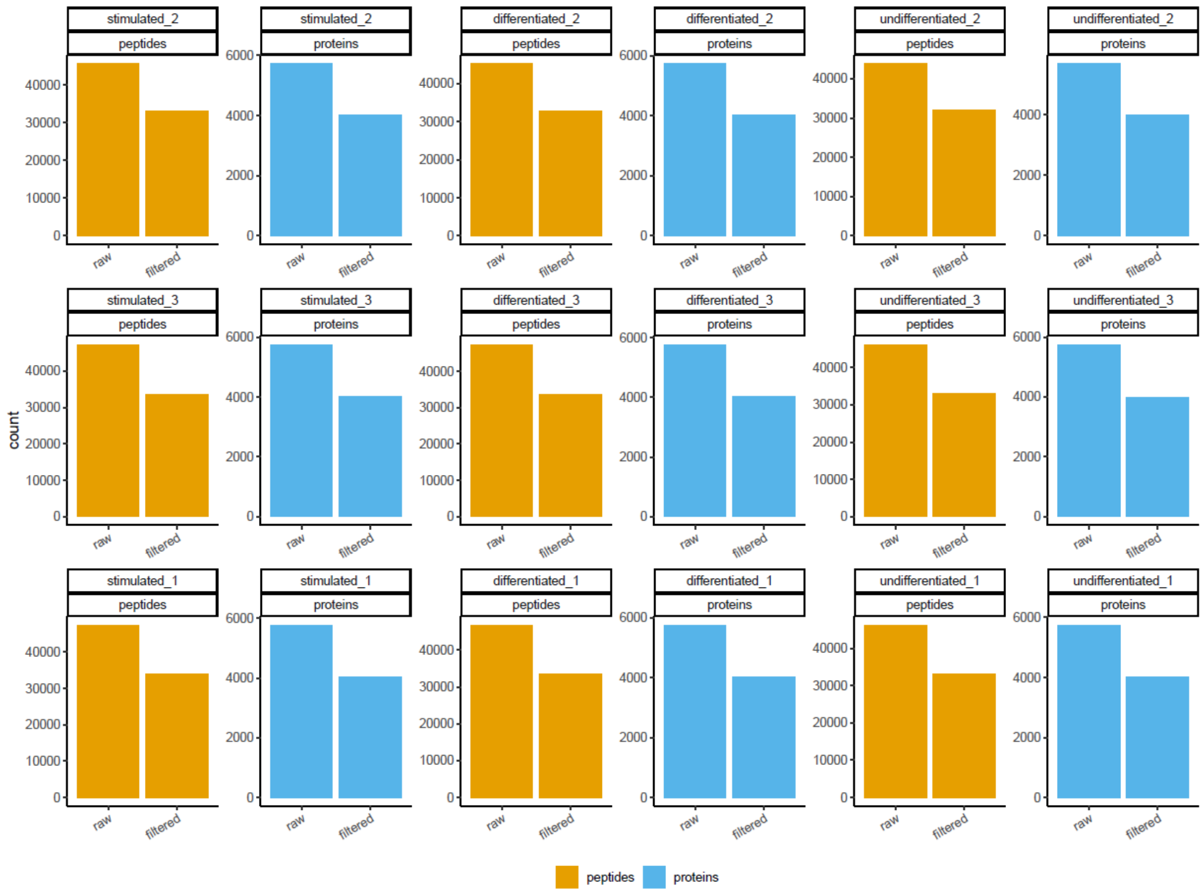


**Supplementary figure 8 – Peptide detection rates and effect of filtering**

Number of peptides (orange) or proteins (in blue) before and after the 3 peptide-filtering steps. The filtering steps are based on the identification of peptides in consecutive fractions and based on the correlation of sibling peptides mapping to the same proteins.

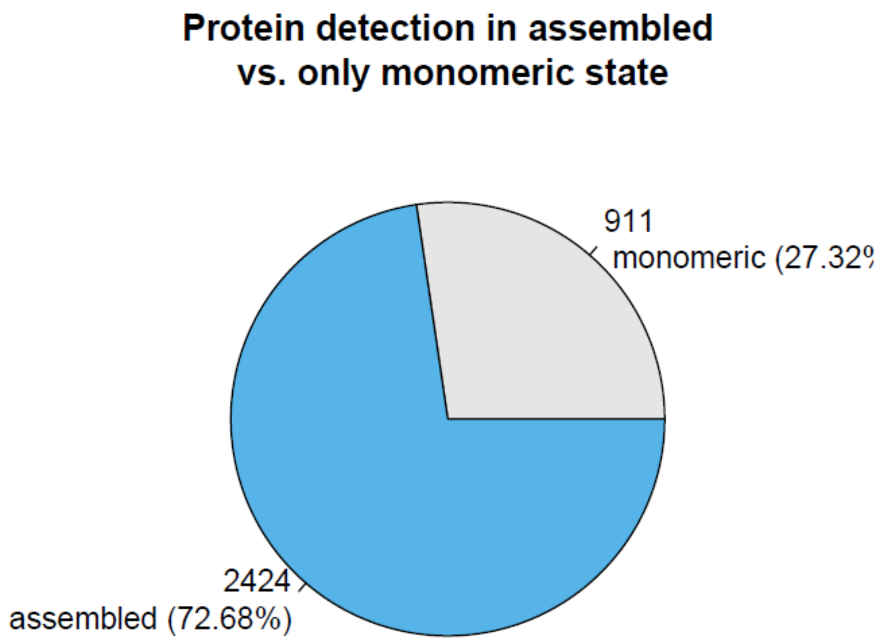


**Supplementary figure 9 – Proteins detected in assembled vs. monomeric state**
The number of proteins for which at least one feature was detected in an 'assembled' state (> 2 x monomer molecular weight) versus those detected only in the 'monomeric' state (< 2 x monomeric molecular weight) are depicted.

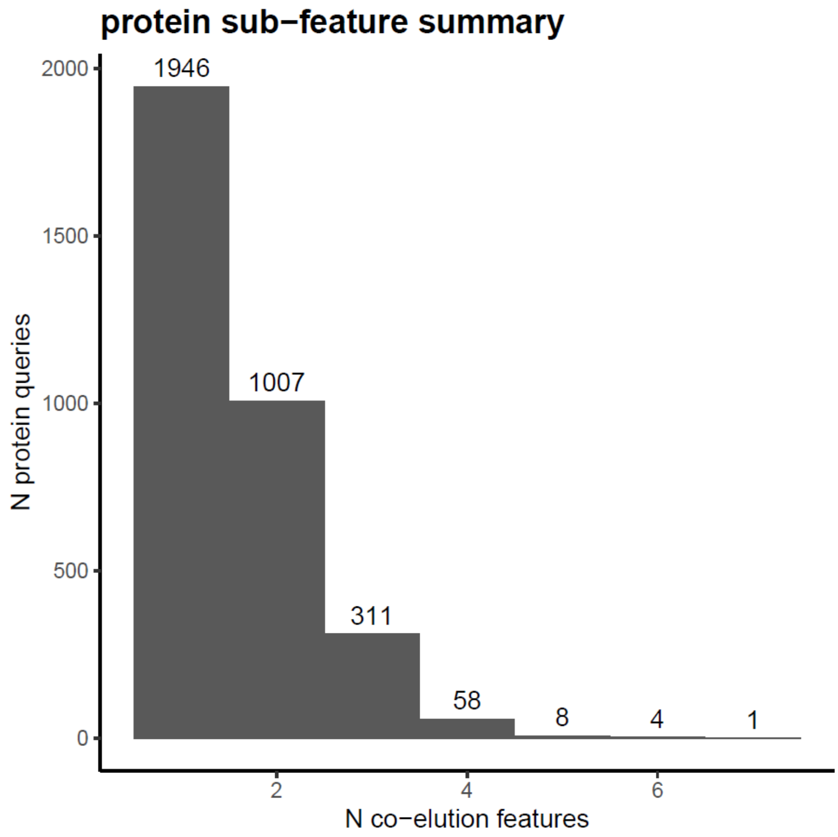


**Supplementary figure 10 – Detected protein feature summary**
The number of SEC elution features a given protein was detected is summarized for all detected proteins after filtering.

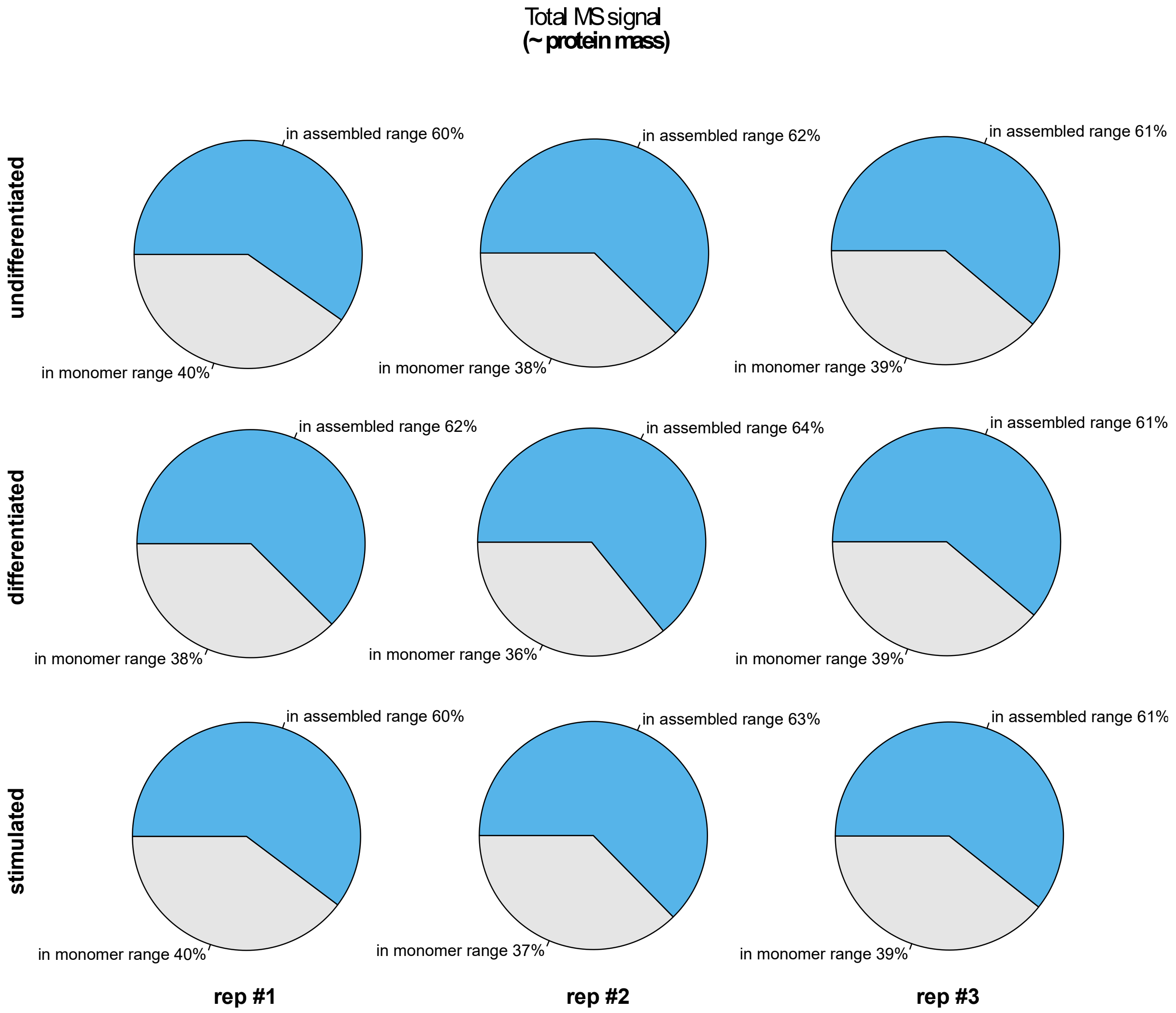


**Supplementary figure 11 – Protein mass in assembled state**

The fraction of protein mass as estimated by the MS signal intensity to be in an assembled (> 2 x monomer molecular weight -- blue) versus monomeric (< 2 x monomer molecular weight -- grey) state is shown.

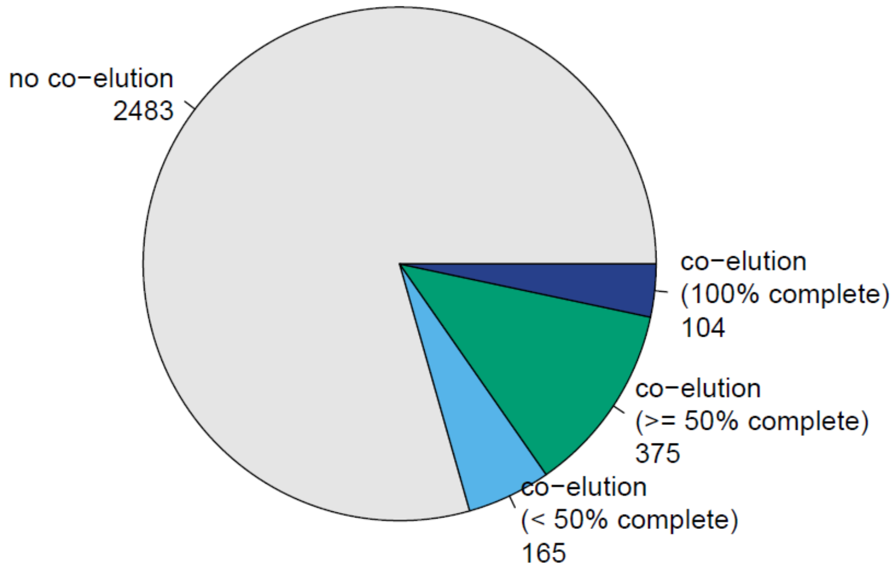


**Supplementary figure 12 – Protein complex detection rates as a fraction of complex hypotheses**
The number and fraction of protein complexes detected in THP1 at 5% FDR at the complex-detection level from the merged CORUM and String (clustered) hypothesis set are shown.

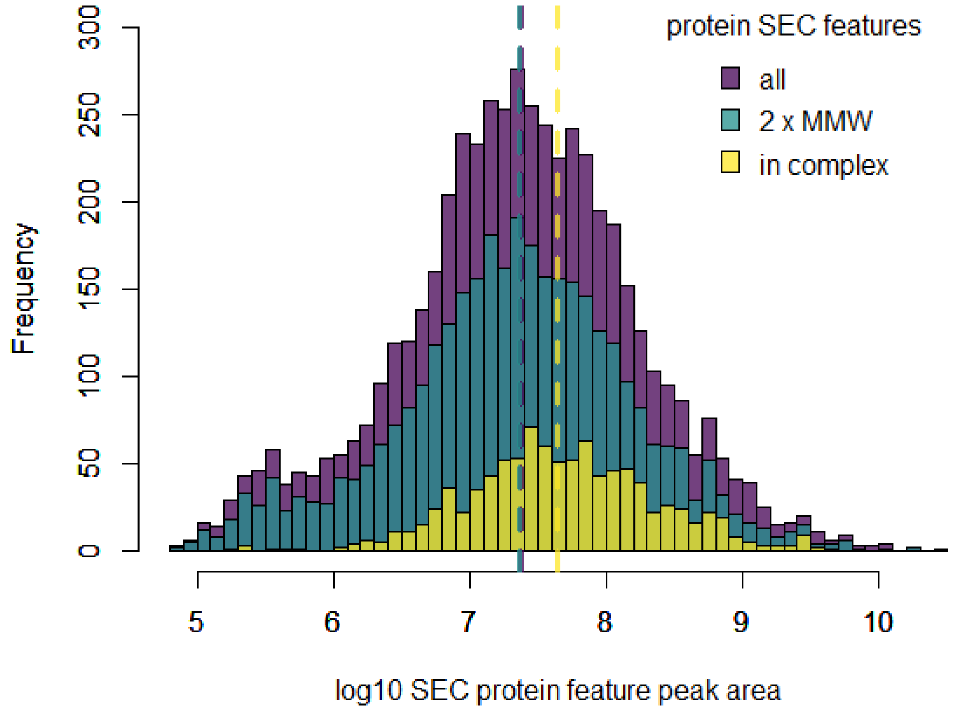


**Supplementary figure 13 – Distribution of SEC protein features dependent on MW and detection in complex-centric analysis**

The distribution of all SEC protein features (purple), the subset of those features that appear at >2x the expected monomer weight and are expected to be in complex (green), and those that are detected as a subunit of protein complex in the complex-centric analysis (yellow) are shown. Matching of protein SEC features to protein complexes is anti-conservative using an apex fraction tolerance of ±5 SEC fractions.
